# A novel subtype of reactive astrocytes critical for HIV associated pain pathogenesis

**DOI:** 10.1101/2022.08.03.502665

**Authors:** Junying Zheng, Michael Spurgat, Shao-Jun Tang

## Abstract

Pathological pain is common in HIV patients, but the underlying mechanism remains elusive and therapeutic targets for effective treatment have not been identified. Reactive astrocytes are specifically activated in the spinal dorsal horn (SDH) of HIV patients with pathological pain and required for the development of HIV-associated pain in mouse models. These findings suggest a key role of reactive astrocytes in HIV-associated pain pathogenesis. However, due to the heterogeneity of reactive astrocytes, the pathogenic subtype is unknown. Using single-nucleus RNA-seq (snRNA-seq) transcriptomic analysis, we identified a novel subtype of HIV-pain associated astrocytes (HIPAs) in the lumbar spinal cord of the HIV -1 gp120 transgenic model. HIPAs were galectin 3 (Gal3)-positive and had transcriptomic signatures of phagocytosis and inflammation; they were also induced in the spinal cord of HIV patients. We showed HIPAs phagocytosed neuronal and synaptic components and were associated with neuronal degeneration. We found that knockout (KO) of Gal3 in gp120 transgenic mice severely diminished HIPAs. Interestingly, the activation of other astrocytes (e.g., homeostatic astrocytes) were also diminished in the Gla3 KO/gp120 transgenic mice. These results indicate that Gal3 is critical for gp120 to induce HIPAs, and that Gal3 may directly or via HIPAs to control the activation of other subtypes of astrocytes. Finally, we showed that the loss of HIPAs caused by Gal3 KO was associated with attenuated neuronal degeneration, neuroinflammation, and pain in gp120 transgenic mice. Collectively, our data suggest that HIPAs are a Gal3-expressing astrocytic subtype that mediates gp120-induced neurodegeneration and neuroinflammation in the spinal pain neural circuit during pain pathogenesis and is a potential cell target for treating HIV-associated pain.

## Introduction

Pathological pain is a common comorbidity of HIV patients, but the underlying mechanism is unclear^1^. Among the neuropathology in the pain neural circuits of HIV patients, astrogliosis (reactive astrocytes) in the spinal cord dorsal horn (SDH) is particularly evident^2^. Importantly, reactive astrocytes were observed in the SDH of only the HIV patients who developed neuropathic pain but not in the SDH of the patients without pain disorders^2^. These findings from studies on human spinal specimens indicate that reactive astrocytes play a critical role in the pathogenesis of HIV-associated pain.

Under physiological conditions, spinal astrocytes modulate pain processing in neuronal circuits by maintaining circuitry homeostasis via their diverse biological activities such as neurotransmitter uptake and release of gliotransmitter^3^. When the homeostatic function is disturbed in reactive astrocytes, their associated pain neuronal circuits may become sensitized for activation. Reactive astrocytes in the SDH have been reported in animal models of various types of pathological pain^4,5^, including the models of HIV-associated pain^6^. In support of their critical role in pain pathogenesis, ablation of reactive astrocytes blocks the expression of mechanical allodynia induced by HIV-1 gp120^7^. Concomitantly, the astrocytic ablation also reverses gp120-induced neural circuit polarization (NCP) in the SDH^7^. Similarly, ablation of reactive astrocytes also blocks opioid-induced hyperalgesia and its associated NCP in the SDH^8^.

Astrocytes are functionally, morphologically, and spatiotemporally heterogenous across CNS regions^9^. During astrogliosis, a spectrum of subtypes of reactive astrocytes are expected to generate, including the better characterized A1 and A2 subtypes^10^. More recently, snRNA-seq analyses have identified distinct astrocyte subtypes associated with different diseases^11^. These astrocyte subtypes have characteristic gene expression signatures that are implicated in specific cellular pathways such as extracellular matrix remodeling^12^, plaque degradation^13^, inflammation, and anti-inflammation^14^ and may contribute to the relevant pathogenesis. However, despite the emerging consensus about the critical role of reactive astrocytes in pain pathogenesis, little is known about the specific pain-pathogenic astrocytic subtypes and how they might contribute to the underlying pathogenesis.

In this study, we performed snRNA-seq analysis of the lumbar spinal cord of HIV-1 gp120 transgenic (gp120Tg) mice^15^, a model of HIV-associated pain, and identified a novel subtype of astrocytes. This astrocytic subtype has the transcriptomic signatures of phagocytosis and inflammation and expresses galectin 3 (Gal3). We show that they are phagocytes of neurons and synapses. Interestingly, Kockout (KO) of Gal3 leads to depletion of this subtype of astrocytes and blockage of the development of mechanical allodynia in the gp120Tg mice, suggesting their association with the pathogenesis of gp120-inudced pain. We hence name this subtype of astrocytes as HIV pain-associated astrocytes (HIPAs). We further show that depletion of HIPAs by Gal3 KO also significantly attenuated neuronal degeneration and neuroinflammation in the spinal cord of gp120Tg mice. These findings indicate that HIPAs may contribute to gp120-induced pain by promoting neurodegeneration and neuroinflammation in the spinal pain neural circuits.

## Results

### Identification of the Gal3+ astrocytes by snRNA-seq transcriptomic analysis

To investigate the cellular impact of gp120 in the spinal cords, we performed snRNA-seq transcriptomic analysis of spinal cords of the Gfap-gp120 transgenic (gp120Tg) mice (4-5 months)^15^, which develop mechanical allodynia (**S. Fig. 1**). Single nuclei were prepared from lumbar regions of the cord and used for cDNA synthesis using 10X Genomics (10X Chromium v3 chemistry). UMAP clustering of 20284 single nuclei dissociated from three of wild-type (WT) and three of gp120Tg mice generated a detailed atlas of transcriptionally distinct populations of spinal cells, including different types of neurons and glia (**Fig. 1a**).

**Figure 1.**
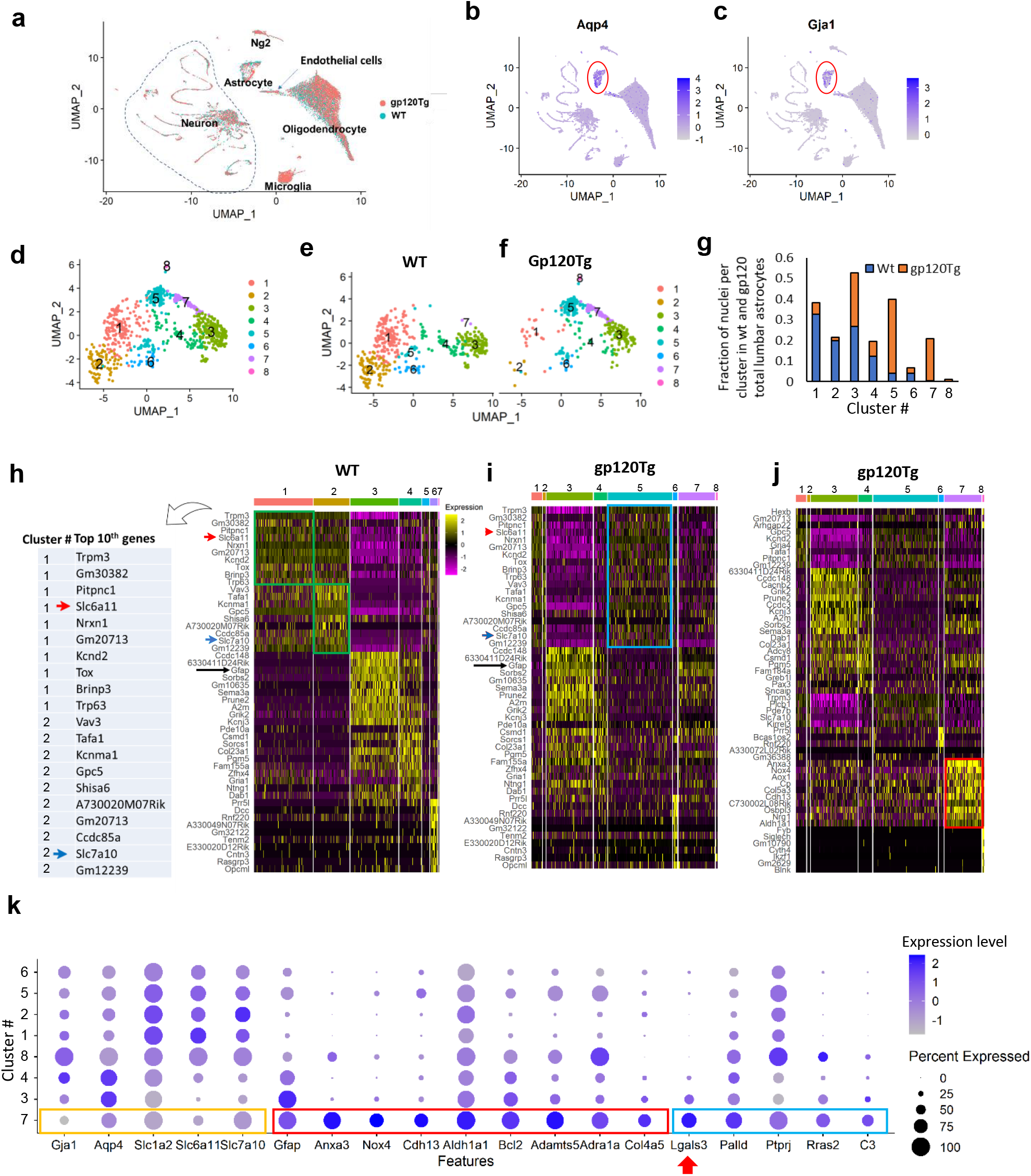
Single-nucleus RNA-seq (snRNA-seq) identified a novel astrocytic subtype in the lumbar spinal cord of gp120Tg mice. **a.** An overview of cell clusters. Major spinal cell types revealed by integrated clustering analysis (by UMAP) of the snRNA-seq datasets from WT and gp120Tg lumbar spinal cords. **b and c.** Identification of astrocytes. The astrocytic cluster (red circle) identified by the feature plots of astrocyte marker genes Aqp4 (b) and Gja1 (c). **d.** Astrocytic subtypes. The snRNA-seq data of astrocytes from WT and gp120Tg mice were subset for re-clustering. Eight astrocytic subtypes were identified. **e** and **f.** Differential astrocytic subtypes in WT and gp120Tg spinal cords. Splitting the merged astrocyte clusters (d) based on their genotypes revealed marked differences in specific subtypes from WT (e) and gp120Tg (f) mice; note the marked decrease of clusters 1 and 2 and increase of cluster 5. Cluster 7 was a subtype specifically in gp120Tg mice. **g.** Quantitation of individual subtypes in WT and gp120Tg mice. Shown here were fractions of nuclei of individual clusters in the total WT or total gp120Tg lumbar spinal astrocytes, respectively. Note that clusters 1 and 2, the two major clusters in the WT were dramatically decreased whereas cluster 5 was increased in gp120Tg mice. Cluster 7 was present in gp120Tg but not WT spinal cords. **h**. Cluster-specific markers in the WT astrocytes. Heatmap shows the expression of top 10 cluster-specific marker genes in astrocytes from the WT mice. Each column represented a nucleus, each row represented a gene. The markers in clusters 1 and 2 (green rectangles) were exemplified in the list on left. Note cluster 1 and 2 express high level of astrocyte transporter genes Slc6a11 (red arrow) and Slc7a10 (blue arrow) and very low level of Gfap (black arrow). On the contrary, Cluster 3 and 4 express very low level of Slc6a11 and Slc7a10 but high level of Gfap. **i.** Expression of the WT cluster-specific markers (h) in the gp120Tg astrocytes. Note cluster 5 in the gp120Tg mice expressed the marker genes of both cluster 1 and cluster 2 from the WT mice. Cluster 3 and 4 from the WT and gp120Tg mice express the same set of markers. **j**. Cluster-specific markers in gp120Tg astrocytes. Heatmap showed the expression of the top10 cluster-specific markers in astrocytes from the gp120Tg mice. Note cluster 7 expressed a unique set of gene markers (red rectangle). **k**. Expression profiles of selected genes in different astrocyte subtypes shown by a dot plot. Cluster 7 has low levels of expression of genes related to astrocytic homeostatic function (orange rectangle), but high levels of expression of genes related to inflammation (red rectangle) and phagocytosis (blue rectangle). scale bar: Log value. Red arrow points to the expression profiles of Lgals3 (gene encoding Gal3). Note the selective expression of Lgal3 in cluster 7.

Our prior studies revealed an association of reactive astrocytes in the spinal dorsal horn (SDH) with the development of neuropathic pain in HIV patients^2^. Using a gp120 model mouse, we demonstrated the critical role of reactive astrocytes in the pathogenesis of HIV-associated pain^7^. To identify gp120-induced specific subtypes of astrocytes contributing to pain pathogenesis, we subset clusters of astrocytes revealed by the feature plots of astrocytic marker genes Aqp4 and Gja1 (**Fig.1b, 1c**). These pooled astrocytes were subjected for further dimension reduction analysis^16^. We identified eight astrocytic subclusters from the combined WT and gp120Tg dataset (**Fig. 1d**). Splitting the clusters based on their genotypes (WT vs. gp120Tg) revealed an evident shift of between WT and gp120Tg astrocytic clusters (**Fig. 1e** and **1f**). Compared with their WT counterparts, the gp120Tg astrocytes in clusters 1 and 2 were markedly decreased, while the gp120Tg astrocytes in cluster 5 were increased. Cluster 7 was specifically identified in the gp120Tg mice (**Fig.1e-1g**).

These astrocytic clusters can be distinguished by cluster-specific marker genes (identified using Seurat FindAllMarkers from the WT and gp120Tg datasets respectively) (**S. Table 1; S. Table 2**). Cluster-specific markers are the upregulated and downregulated genes (with adjusted p-value<0.05) of the single astrocytic cluster compared to all other astrocytic clusters. Figure 1h showed the expression of the top 10 cluster-specific marker genes in the WT dataset. The WT astrocytic clusters were separated into two major categories based on their Gfap expression levels, with clusters 1, 2, 5, 6 manifesting low (Gfap^low^) and clusters 3 and 4 manifesting high Gfap (Gfap^high^) expression (**Fig.1h**; **S. Fig.2a**); these results confirmed previous observations^17^. Clusters 1 and 2 were the two major subpopulations expressing Slc6a11 and Slc7a10 (**Fig.1h**). Slc6a11 encodes an astrocyte neurotransmitter transporter playing key roles in GABA uptake^18^. Slc7a10 encodes a sodium-independent amino acid transporter with a primary role related to modulation of excitatory glutamatergic neurotransmission^19^. Clusters 3 and 4 expressed high Gfap level but low level of Slc6a11 and Slc7a10 (**Fig.1h**).

The astrocytic clusters in the gp120Tg mice manifested a clear shift from Gfap^low^ toward Gfap^high^ populations (**S**. **Fig.2b**). To assess the biological relation between the WT and gp120Tg cells in the same cluster, we examined the expression profiles of the top 10 cluster-specific marker genes of the WT dataset in the gp120Tg dataset (**Fig. 1i**). We found that clusters 3 and 4 from the gp120Tg expressed the same set of marker genes of WT cluster 3 and 4. Further, the fraction of these clusters in the total spinal astrocytes did not change markedly between the WT and gp120Tg mice (**Fig.1g**). These data together suggested that these subtypes of astrocytes were not drastically altered in the gp120Tg mice. In contrast, cluster 5 was drastically increased while clusters 1 and 2 were decreased in the transgenic mice. Cluster 5 from the gp120Tg mice expressed the marker genes of WT cluster 1 and 2 (e.g., Slc6a11, Slc7a10) (**Fig.1h, 1i, Fig. 6b, 6f**). The shift of astrocytic clusters expressing these homeostatic genes toward cluster 5 suggested an activation of homeostatic astrocytes in the gp120Tg mice.

**Figure 2.**
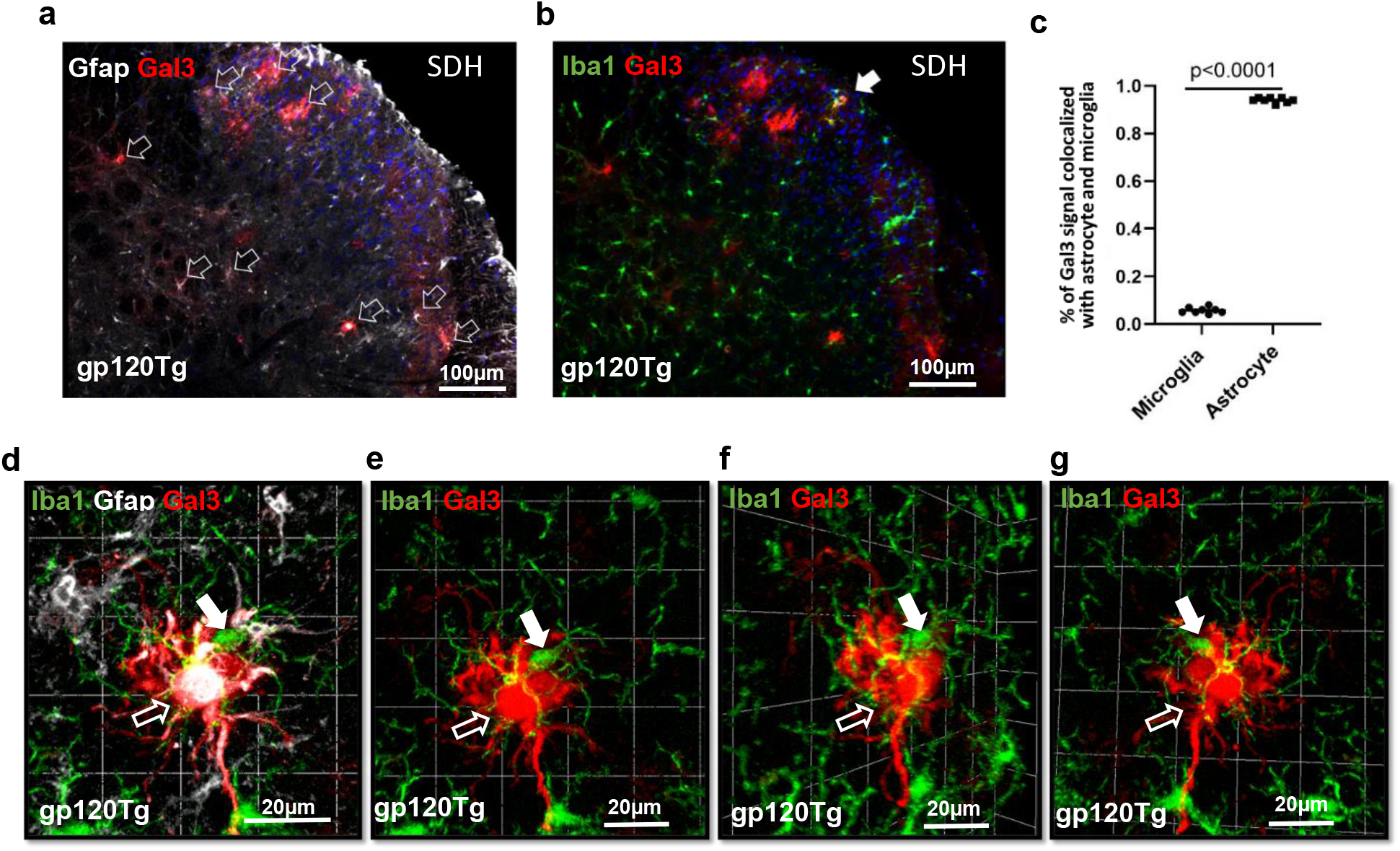
Gal3 was predominantly expressed in spinal astrocytes rather than microglia in gp120Tg mice. **a** and **b** Double immunofluorescent staining (IF) of Gal3 (red) with astrocytic marker (Gfap) (white) and microglial marker Iba1 (green) respectively showed most Gal3^+^ cells were also GFAP^+^ (white open arrows) (a) in the gp120Tg lumbar spinal cord, although Gal3^+^ microglia were occasionally observed (white filled arrowhead) (b). SDH: spinal dorsal horn. **c,** Quantitative summary of Gal3^+^ microglia and astrocytes. **d,** A 3D confocal image of a cell clump of microglia (filled arrow) and an astrocyte (open arrow) with Gal3 signals (red). **e-g**: Different optical sections of the clump showed that Gal3 was exclusively in the astrocyte (open arrow) but not in the nearby microglia (filled arrow).

Cluster 7 is a novel astrocyte subtype that was exclusively identified in the gp120Tg mice (**Fig.1f, 1g**). We named this cluster as <u>HI</u>V-<u>p</u>ain associated <u>a</u>strocytes (HIPA) based on their potential pathogenic contributions. HIPAs comprised of 20% of total gp120Tg spinal astrocytes. Gene expression heatmap of the top 10 cluster-specific marker genes (**S. Table 2**) from the gp120Tg dataset suggested a unique set of genes expressed in HIPAs (**Fig.1j**) including Anxa3, Nox4, Aox1, Cp, etc. To gain insight into their potential biological function, we imported the 112 HIPA (cluster 7)-specific marker genes from the gp120Tg dataset (**S. Table 2, cluster 7**) into Ingenuity Pathway Analysis (IPA). IPA identified 33 well-characterized inflammatory genes (**S. Fig.3**) and 7 well-characterized phagocytotic genes (**S. Fig.4**) which predicted an increased biological activity in inflammation and phagocytosis (**S. Fig.3** and **S. Fig.4**), respectively.

To visualize the expression profiles of the top IPA-selected genes related to astrocytic homeostasis, inflammation, and phagocytosis, we generated a dot plot using the combined WT and gp120Tg dataset (**Fig. 1k**). The dot plot showed that HIPAs (cluster 7) expressed low levels of homeostatic genes (Gja1, Aqp2, Slc1a2, etc.) but high levels of inflammatory genes (Gfap, Anxa3, Nox4, Adamts5, Cdh13, Aldh1a1, Bcl2, Adra1a, Col4a5) and phagocytotic genes (Lgals3, Palld, Ptprj, Rsas2, C3) (**Fig.1k**). These data indicate HIPAs are both inflammatory cells and phagocytes.

### Features of HIPAs

Among the phagocytosis-or inflammation-related up-regulated genes in HIPAs, Lgals3 was predominantly expressed in this novel astrocytic subtype, compared to others (**Fig. 1k**). Importantly, in the context of pain pathogenesis, Gal3 was reported to play a critical role in neuropathic pain in a peripheral nerve injury model^20,21^. Hence, we were interested in validating, at a protein level, Gal3 is a marker for HIPAs. To this end, we performed double fluorescent immunostaining of Gfap and Gal3 and confocal imaging. As Gal3 was reported to predominantly expressed in microglia in the brain^22^, we also performed a control experiment of co-staining of Gal3 and microglial marker Iba1. We observed that Gal3 was almost exclusively expressed in astrocytes but not in microglia in the gp120Tg spinal cords (**Fig. 2a, 2b**); about 95% of the Gal3^+^ cells were also Gfap^+^ (**Fig. 2c**). When Gal3^+^ cells were in a cluster containing both Gfap^+^ and Iba1^+^ cells (**Fig. 2d**), images of optical sections showed Gal3 signals were in astrocytes but not microglia (**Fig. 2e~2g**). These data confirm that, as suggested by the snRNA-seq data, Gal3 is a specific marker for HIPA in spinal cords.

### Spatial distribution of HIPAs in the spinal cord

To confirm the induction of HIPAs in gp120Tg spinal cords, we performed western blotting (**Fig. 3a**) and immunostaining analysis (**Fig. 3b-3d**) of Gal3 expression in the spinal cord of WT and gp120Tg mice. We observed that Gal3 level was low in WT but significantly upregulated in gp120Tg spinal cords (**Fig. 3a**), indicating the formation of HIPAs in gp120Tg spinal cords. Immunostaining of Gal3 revealed Gal3^+^ cells were barely detected in WT spinal cords except in the central canal area (**Fig. 3b**) with sparse Gal3^+^ cells detected in marginal areas and dorsal horns (**Fig.3b**). In contrast, Gal3^+^ cells significantly increased in the gp120Tg cord, especially in the grey matter (**Fig. 3c, 3d**). These observations confirm the induction of Gal3^+^ HIPAs in the spinal cord of gp120Tg mice.

To gain insight into the origin of HIPAs, we compared the spatial distribution patterns of Gal3^+^ cells in the WT and Tg spinal cords. We found that in WT mice Gal3^+^ cells were mainly restricted around the central canal of the spinal cord (**Fig.3b, 3e, 3h**). Figure 3e~3f are high-power magnification images showing Gal3^+^ cells lined in layers (**Fig. 3e**) in the central canal of WT mice, and they did not express astrocytic marker protein Gfap (**Fig.3f, 3g**). The high-power magnification images further confirmed there was no Gal3^+^Gfap^+^ (HIPAs) cells outside the central canal (**Fig.3h**) of the WT mice. In contrast, in the gp120Tg mice, Gal3^+^ cells appeared to spread from the central canal regions to spinal dorsal and ventral horns, forming a clear trajectory starting at the central canal region (**Fig. 3d**). Meanwhile, Gal3 signals in the central canal were diminished in the gp120Tg mice (**Fig. 3i**). Figure 3l show Gal3 signal migrated out of the central canal in Gal3^+^Gfap^+^ (HIPAs) cells in the gp120 mice. In contrast to the Gal3^+^ cells in WT central canal, cells in central canal of the gp120 mice expressed astrocytic marker Gfap (**Fig.3j, 3k**). Collectively, these spatial distribution patterns of Gal3^+^ cells are consistent with the idea that Gal3^+^ cells of the central canal are induced to migrate out to differentiate into HIPAs in the spinal dorsal and ventral horns. The expression of Gfap in the gp120Tg central canal cells suggested the central canal cells showed astrocytic lineages in the gp120 mice.

**Figure 3.**
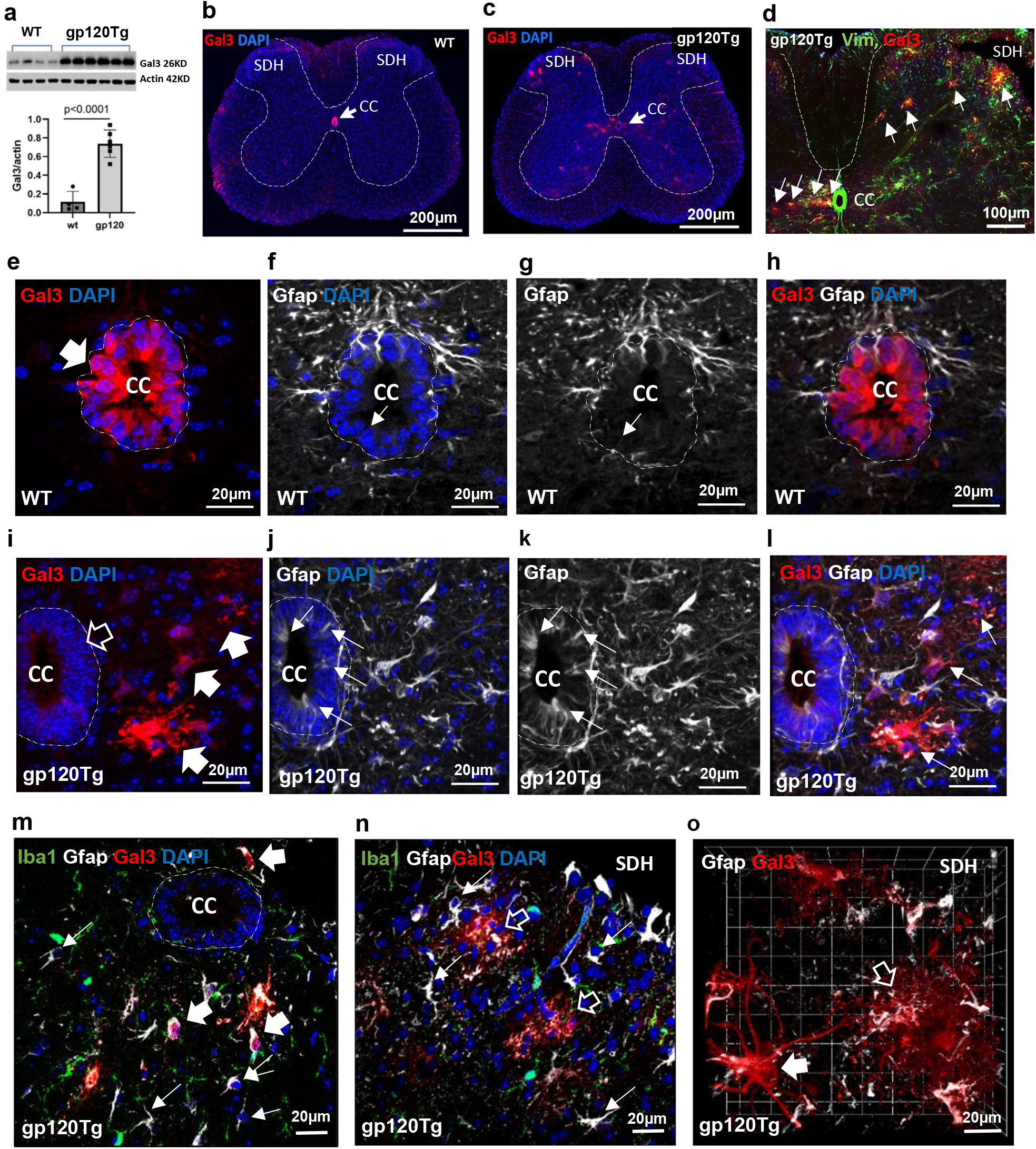
Induction and morphological features of HIPAs in the spinal cord of gp120Tg mice. **a.** HIPA marker Gal3 was significantly upregulated in the spinal cords of gp120Tg mice. **b.** IF revealed Gal3^+^ cells were predominantly at the central canal (CC) region (arrow) in WT mice. **c.** The spatial distribution of HIPAs in the gp120Tg spinal cord shown in a low power image. HIPAs were scattered in the grey matter. Compared with WT (Fig. 3b), Gal3 signals around the central canal (arrow) was diminished. Dash lines enclosed the gray matters. **d**. An IF image of HIPAs (white arrows) that form a trajectory from the CC region to spinal dorsal horn (SDH) of the gp120Tg mice. Dash lines enclosed the gray matters. **e-l**, The distribution of Gal3 (red) protein in spinal CCs (encircled by dash lines) of WT and gp120Tg mice. e, In the WT mice, Gal3 protein was strictly and strongly distributed in Gal3^+^ cells lined in the CC (wide filled arrow); f-h, Gal3^+^ cells in the WT CC did not expressed astrocytic marker Gfap; h, no Gal3^+^Gfap^+^ astrocytes (HIPAs) were detected outside the WT CC. i, In the gp120Tg mice, the expression of Gal3 in CC was severely diminished (wide open arrow), instead strong Gal3 protein were expressed outside CC (wide filled arrows); j and k, Compared to WT mice (f, g), cells in the gp120Tg CC expressed Gfap (thin arrows); l, HIPAS were induced outside the gp120Tg CC (thin arrows). **m-o,** Confocal images showing the heterogenous morphology of HIPAs in different locations in the gp120Tg spinal cord. m, Most HIPAs (wide filled arrows) near the CC region (encircled by dash line) were small and had Gal3 protein concentrated in the nucleus area. n, In the SDH, HIPAs were larger in size with various shapes and Gal3 was distributed throughout the whole astrocyte including the processes (open arrows); m and n, Gal3-negative astrocytes (thin arrows) were smaller in size with fewer short processes. **o**. A stacked confocal image to show the various morphologies of large HIPAs in SDH. Filled arrow pointed to a HIPA with a relative smaller volume and fewer and long processes. Open arrow pointed to a HIPA with bigger cellular body and many short processes. Note the expression of Gal3 (red) across the whole cells in these HIPAs.

HIPAs manifested morphological alterations during this process. In areas nearby the central canal in gp120Tg mice, most HIPAs were relatively small with shorter processes (**Fig. 3m**). At this stage, Gal3 protein was mainly in nuclei area (**Fig.3m**). HIPAs in regions away from the central canal regions, especially in the dorsal horn, showed enormous cell volume and more processes (**Fig. 3n, 3o**). At this stage, Gal3 protein was observed across the whole cells (**Fig.3o**) and these HIPAs showed various morphologies. Figure 3o is a stacked confocal image to show enormous HIPAs with different morphologies in SDH; one with relative smaller cell body and long processes and the other with big swollen cellular body and many short processes. Overall, during their differentiation in the spinal cord of gp120 Tg transgenic mice, HIPAs increased their cell complexity and changed the intracellular distribution pattern of Gal3.

To evaluate the relevance of HIPAs induced in the gp120Tg model, we performed analysis on spinal autopsies from HIV patients. We found Gal3 protein was significantly induced the spinal cord of HIV patients, compared with control patients died from non-HIV causes such as endocarditis, pulmonary embolus, and pneumonia (**S. Fig. 5a**). In addition, immunofluorescent staining showed that the number of Gal3^+^ astrocytes was drastically increased in the HIV patients (**S**. **Fig. 5b, 5c**). These data show that, similar to the gp120 Tg model, Gal3^+^ astrocytes are also induced after HIV infection.

### HIPAs are phagocytes of neurons

Prior studies reveal that Gal3-expression cells, e.g., Gal3^+^ macrophages, function as phagocytes^23^. Gal3 phagocytes can release Gal3 as opsonin to label target cells for phagocytosis^24^. Our findings of upregulation of phagocytosis-related genes in HIPAs (**Fig. 1k**) suggest this subtype of Gal3^+^ astrocytes may also have phagocytic activity. To test this hypothesis, we performed experiments to examine if HIPAs phagocytose neuronal components. Because our previous work suggested synaptic degeneration in the SDH of HIV patients with neuropathic pain^2^ and in the spinal cord of a mouse model of HIV-associated pain^6^, we postulated that HIPAs may contribute to the neurodegeneration as phagocytes. By double fluorescent immunostaining of NeuN (a marker of neuronal soma) and Gal3, we identified HIPAs that engulfed NeuN compartments in the SDH of gp120Tg mice (**Fig. 4a-4g**).

**Figure 4.**
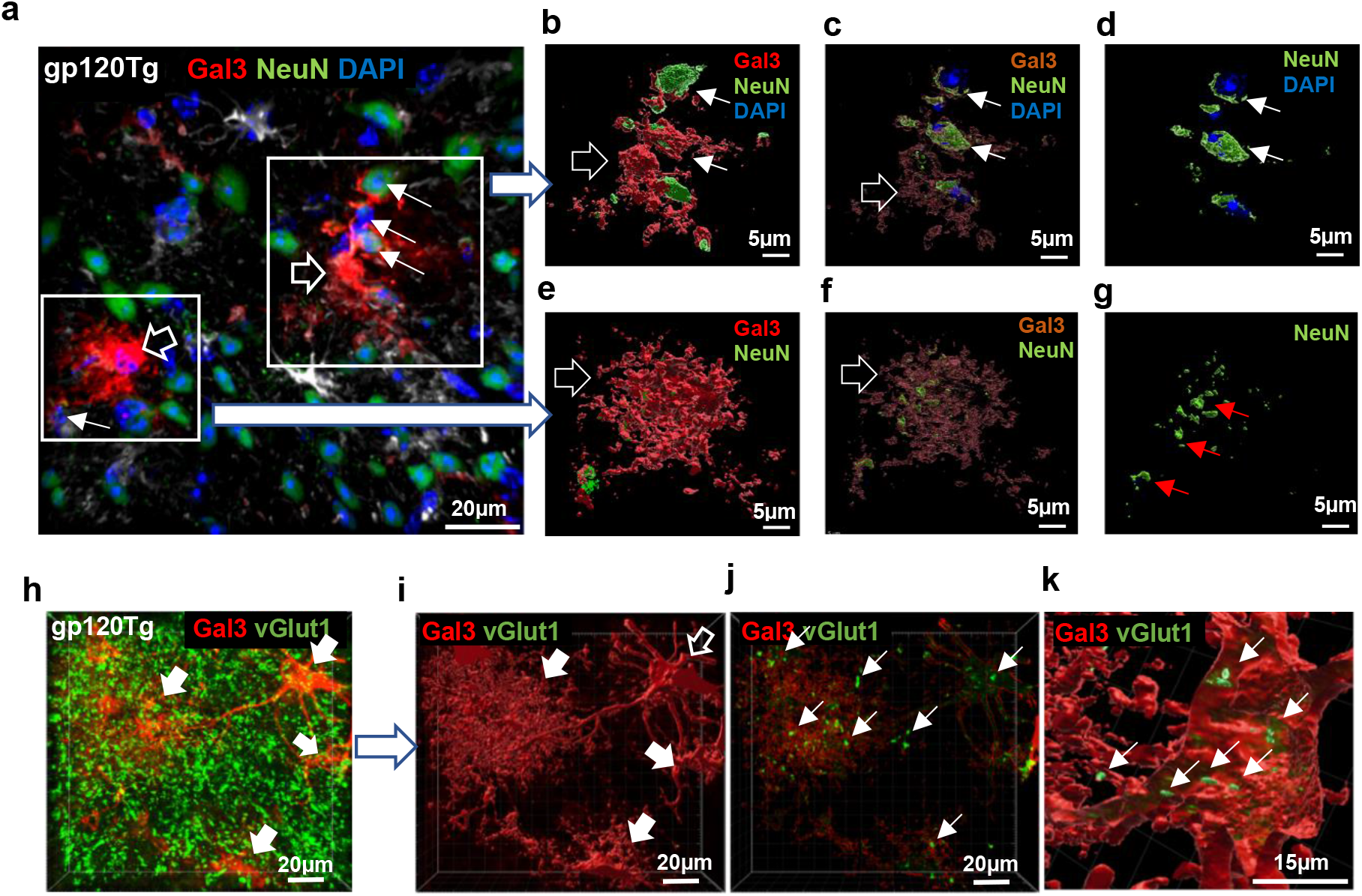
HIPAs engulfed neurons and synapses in the gp120Tg spinal cord. **a.** Confocal images of HIPAs (Gal3^+^) and neurons (NeuN^+^) revealed by double immunofluorescent staining. Shown were two representative HIPAs (red) (open wide arrows) in close interactions with neurons (green) (thin arrows). **b-g,** Imaris 3D image segmentation of the selected area in a. This image analysis removed all the signals that were not interact with HIPAs to visualize NeuN signals engulfed by HIPAs. b-d, Imaris image segmentations of one region in (a) to show intact neuronal somas (thin white arrows) engulfed by a HIPA. e-g. Imaris image segmentations of another regions in (a) to show neuronal debris (thin red arrows) inside a HIPA. The engulfed neuronal somas revealed by creating the solid opaque surfaces (open arrow)(b and e), transparent surfaces (open arrow) (c and f), and by removing HIPA components (d and g). **h,** Confocal image of HIPAs (wide filled arrows) and synapses (green dots) in the SDH of gp120Tg mice. **i** and **j.** Imaris segmentation of the image in h to reveal HIPA-synapse interaction. i-k, This image analysis removed all the synapse signals (green dots) outside HIPAs (wide open and filled arrows). HIPAs were revealed by creating the solid opaque surfaces (wide filled and open arrows) (i). j, The engulfed synapses (thin arrows) were viewed by creating transparent surface of HIPAs in i; **k**. The imaris clipping plane tool truncated a 3D HIPA to view the synapses (green) (arrows) inside a representative HIPA (open arrow in i).

Applying Imaris Microscopy Image Analysis Software, we visualized neuronal components engulfed by HIPAs. Figure 4a is a confocal image of HIPAs (Gal3^+^cells) and neurons (NeuN^+^) by double immunofluorescent staining. Figure 4a demonstrated two representative HIPAs in close interactions with neurons in the SDH of gp120Tg mice. Imaris 3D image segmentation analysis removed all the signals that were not interact with HIPAs (**Fig. 4b~4g**) to visualize NeuN signals engulfed by HIPAs. Neuron compartments engulfed by HIPA were revealed by creating the solid opaque surfaces (**Fig. 4b, 4e**), transparent surfaces (**Fig. 4c, 4f**) of HIPAs and by removing all HIPA components (**Fig. 4d, 4g**). These images clearly demonstrated the intact neuronal somas (**Fig. 4d**) and debris (**Fig.4g**) inside HIPAs.

To evaluate the potential involvement of HIPAs in synaptic degeneration, we also performed double immunofluorescent staining of Gal3 and vGlut1 (a synaptic marker). Figure 4h demonstrated the interactions of HIPAs with synapses in the SDH of gp120Tg mice. Imaris segmentation analysis removed all the synapse outside HIPAs (**Fig.4i~4k**). The engulfed synapses were revealed by creating solid opaque surfaces (**Fig. 4i**) and transparent surfaces (**Fig. 4j**) of HIPAs. Applying Imaris clipping plane tool to truncate a 3D HIPA cell, the synapses inside a representative HIPA were clearly viewed (**Fig.4k**). These findings indicate that HIPAs are neuronal phagocytes in the SDH of gp120 Tg mice.

### Gal3 knockout (KO) diminished HIPAs in the gp120Tg spinal cord

Because Gal3 is implicated in cell growth and differentiation among other biological function^25^ and is upregulated by gp120 (**Fig.3a**), we sought to test the role of Gal3 in HIPA formation in the gp120Tg spinal cord. To this end, we generated gp120Tg mice with Gal3 knockout (gp120Tg/Gal3^-/-^ mice), by crossing gp120Tg mice^15^ with mice with a Gal3 targeting mutation. Western blotting analysis showed Gal3 KO significantly attenuated the upregulation of protein expression of Vim and Gfap, markers of reactive astrocytes, in the gp120Tg spinal cord (**S. Fig. 6a-6c**).

Next, we sought to determine specifically the potential role of Gal3 in the formation of the HIPA subtype. To this end, we performed snRNA-seq analysis to determine if HIPAs can still form in gp120Tg/Gal3^-/-^ mice. Integrated UMAP clustering analysis of snRNA-seq data of 48,935 nuclei from twelve of wild-type (n=3), gp120Tg (n=3), gp120Tg/Gal3^-/-^ (n=3), and Gal3^-/-^ (n=3) mice showed all major spinal cell types, including different types of neurons and glia (**Fig. 5a**). Astrocytes (**Fig.5b, 5c**) from all four genotypes were subset for a second-dimension reduction analysis. Astrocytic clusters were first displayed in an integrated UMAP plot (**Fig. 5d**). Splitting the astrocytic clusters based on their genotypes revealed that cluster 7 (HIPAs) that was specifically observed in the gp120Tg mice (compared with WT) (**Fig. 5e-5f**), as shown in **Fig. 1e-1f**. Importantly, this group of astrocytes was largely disappeared in the gp120Tg/Gal3^-/-^ mice (**Fig. 5g**, **5i**). This observation suggested the Gal3 KO prevented HIPA formation in gp120Tg/Gal3^-/-^ mice. In support of the diminish of HIPAs, the top 50 upregulated genes in HIPAs, including genes in inflammatory and phagocytotic pathways were also dramatically attenuated in the gp120Tg/ Gal3^-/-^ mice (**Fig.5j-5l**). Also, astrocytes expressing inflammatory genes (e.g., Anxa3) and phagocytotic gene (e.g., C3) compared to WT (**Fig. 5n, 5r**) were markedly increased in the gp120Tg mice (**Fig.5o, 5s**), primarily in cluster 7 (HIPAs); and the increases were drastically attenuated in the gp120Tg/Gal3^-/-^ mice (**Fig. 5p, 5t**). On the other hand, compared to the WT, Gal3 KO in WT mice did not induce evident changes on the expression of those HIPA genes (**Fig. 5j, 5m**) and the astrocytic clusters (**Fig. 5e, 5h, 5n, 5q, 5r** and **5u**). These data indicate that Gal3, which is induced by gp120 (**Fig. 3a**), specifically controls the formation of HIPAs.

**Figure 5.**
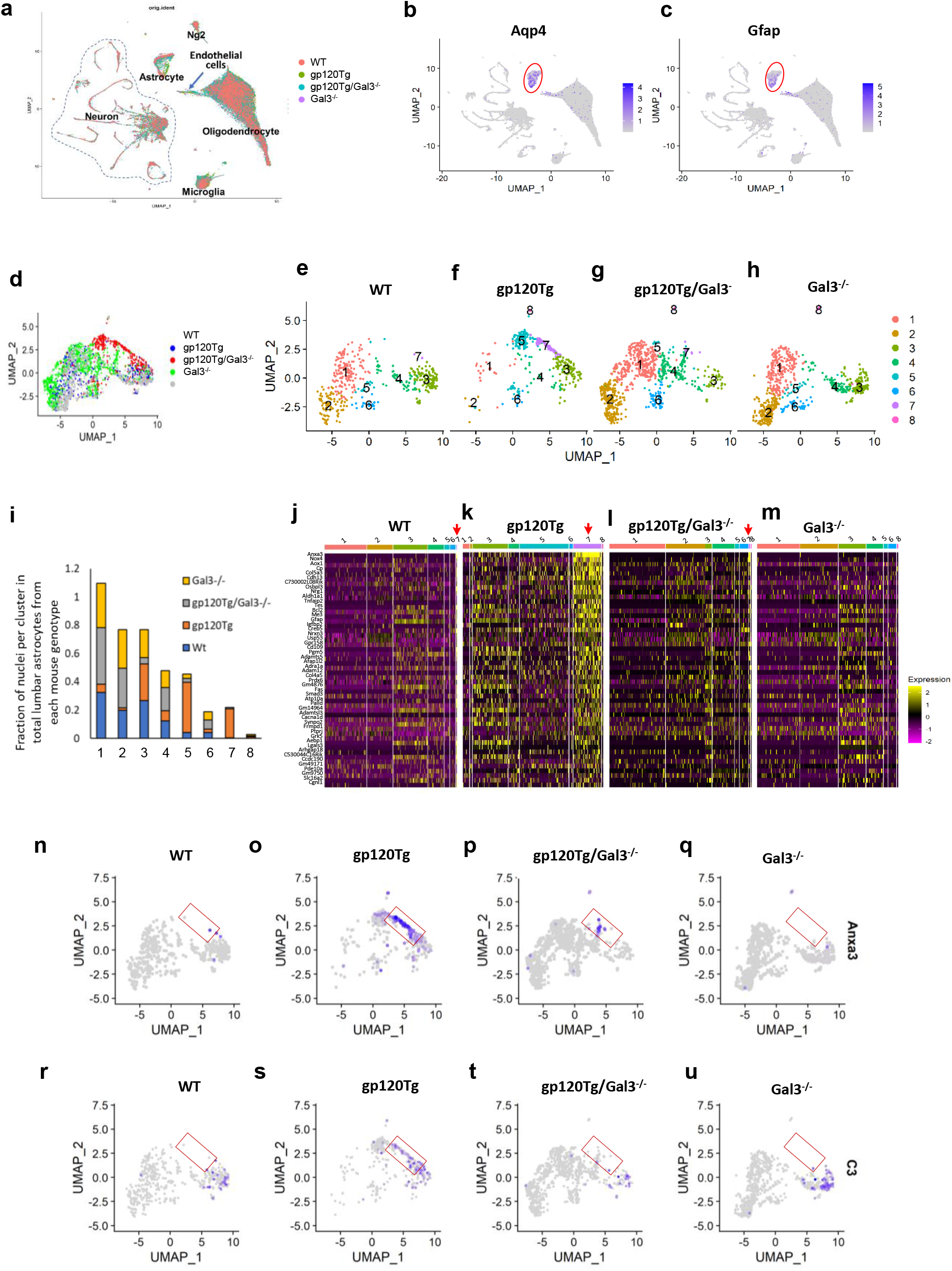
Gal3 KO led to depletion of HIPAs (cluster 7) and attenuation of the activation of other subtypes of astrocytes. **a.** A UMAP cluster plot of the integrated snRNA-seq datasets from WT, gp120Tg, gp120Tg/Gal3^-/-^, and Gal3^-/-^ mice to show the clusters of the main cell types in lumbar spinal cords. **b** and **c**. Feature plots of astrocyte marker genes Aqp4 (b) and Gfap (c) to show the astrocytic cluster (red circles) in the combined UMAP cluster plot. **d.** A UMAP cluster plot of subset analysis of combined astrocytes from of WT, gp120Tg, gp120Tg/Gal3^-/-^, and Gal3^-/-^ lumbar spinal cords. **e-h.** UMAP cluster plots of astrocytes from different genotypes of mice after splitting the overlayed astrocytic clusters (d). Compared with the WT astrocytic clusters (e), there were evident shifts of several gp120Tg clusters (f), including marked decrease of clusters 1 and 2 and increase of cluster 5. Note that cluster 7 was almost exclusively in the gp120Tg plot (f), with only a small number of homologous cells in the counterpart cluster in the WT plot (e). Strikingly, these gp120-induced shifts of astrocyte cluster sizes were reversed by Gal3 KO (g). Compared with clusters in the plot of gp120Tg (f), the clustering pattern of gp120Tg/Gal3 KO astrocytes (g) appeared to shift toward the pattern of WT astrocytes (e), with increased sizes of clusters 1 and 2 and decreased size of cluster 5. Notably, the gp120Tg-specific cluster 7 (i.e., HIPA) was almost completely depleted in gp120Tg/Gal3 KO mice (g). The overall cluster pattern of gp120Tg/Gal3 KO (g) was more similar to that of WT (e) than gp120Tg (f). h, A UMAP cluster plot of astrocytes from Gal3 KO mice. The cluster pattern of Gal3 KO astrocytes (h) was similar to that of WT (e). **i.**Quantitative analysis of the relative population sizes of nuclei of individual clusters from different genotypes. Note that Gal3 KO depleted HIPAs (cluster 7) in gp120Tg mice, and prevented the decrease of the two homeostatic astrocytic subpopulations (cluster 1, 2), and attenuated the increase of cluster 5. **j-m**. Comparison of the expression of the top 50 upregulated genes in HIPA (cluster 7 in f) in gp120Tg (k) with their counterpart genes in WT (j), gp120Tg/Gal3 KO (l), or Gal3 KO astrocytes in heatmaps. These genes included those implicated in inflammatory and phagocytotic pathways. Each row represented a gene, and each column represented a cell. The color bars on top of individual heatmaps represented individual clusters, and the color scale bar on the right represented the expression levels (log2 values). The red arrows point to cluster 7 in individual heatmaps. Note there is no cluster 7 in Gal3 KO mice. Compared with WT (j), these genes were significantly upregulated in cluster 7 of gp120Tg astrocytes (k), but the upregulation was blocked in gp120Tg/Gal3 KO astrocytes (l). Compared with the heatmap of WT astrocytes (j), Gal3 KO only (m) did not evidently alter the expression of these genes. **n-p,**Effect of HIPA depletion on inflammatory astrocytes. Astrocytes expressing inflammatory gene Anxa3 (enclosed by the red rectangles) compared to the WT spinal cord (n) were drastically increased in the gp120Tg cord (o); and the increase was markedly attenuated after HIPA depletion in the gp120Tg/Gal3^-/-^ mice (p). **r-t**, Effect of HIPA depletion on phagocytotic astrocytes. Astrocytes expressing phagocytotic gene C3 (enclosed by the red rectangles) compared to the WT spinal cord (r) were drastically increased in the gp120Tg cord (s), and the increase was markedly attenuated after HIPA depletion (t) in the gp120Tg/Gal3^-/-^ mice. **q** and **u,**Gal3 KO in WT mice did not have noticeable impact on the inflammatory and phagocytotic astrocytes (q, u) compared to that in the WT mice (n, r). Scale of gene expression (log value): blue is high; gray is low.

**Figure 6.**
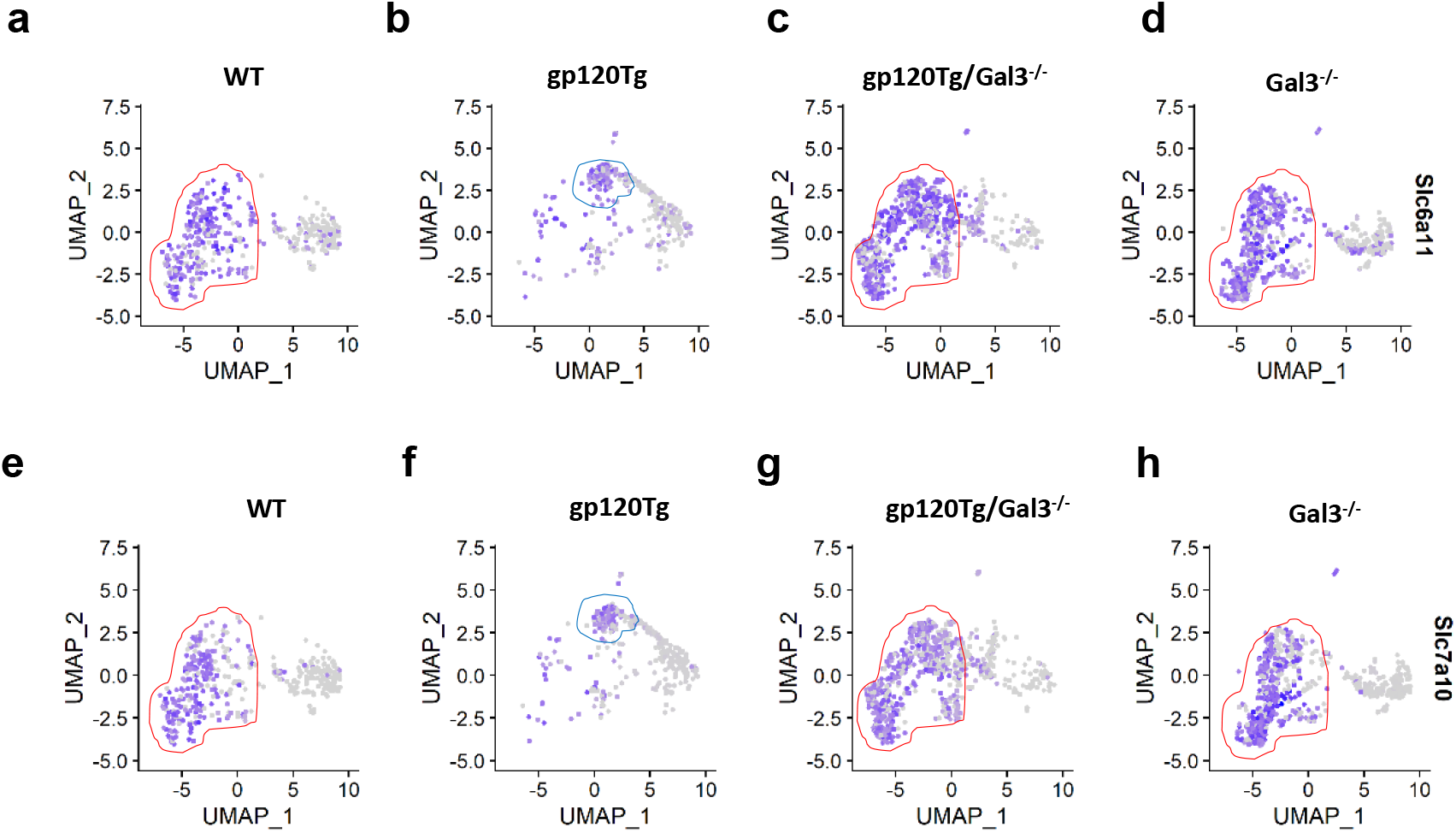
Gal3 KO in the gp120Tg mice attenuated the activation of astrocytes in regulation of neurotransmitter homeostasis. **a-d,** Gal3 KO attenuated the activation of astrocytes related to GABA uptake. Astrocytes expressing Slc6a1, the astrocytic transporter mediating GABA uptake were mainly in cluster 1 and 2 (enclosed by red line) in the WT spinal cord mice (a); shifted to cluster 5 (enclosed by blue line) in the gp120Tg cord which indicating the activation of these astrocytes (b); the shift was prevented by Gal3 KO gp120TgGal3^-/-^ cord (c); Gal3 KO in the WT mice did not have impact on these homeostatic astrocytes (d). **e-h**, Gal3 KO attenuated the activation of astrocytes mediating excitatory glutamatergic neurotransmission.Astrocytes expressing Slc7a10, the astrocytic transporter mediating excitatory glutamatergic neurotransmission were mainly in cluster 1 and 2 (enclosed by red line) in the WT spinal cord mice (e); shifted to cluster 5 (enclosed by blue line) in the gp120Tg cord (f); the shift was prevented by Gal3 KO in the gp120TgGal3^-/-^ cord (g); Gal3 KO in the WT mice did not impact on these homeostatic astrocytes (h). Scale of gene expression (log value): blue is high; gray is low.

### Gal3 KO attenuated the activation of astrocytes in regulation of neurotransmitter homeostasis in the gp120 spinal cord

Gal3 KO in gp120 transgenic mice severely diminished HIPAs. Interestingly, we found the activation of homeostatic astrocytes were also diminished in the gp120Tg/Gal3^-/-^ transgenic mice. Figure 6a~6h showed the effect of Gal3 KO on the activation of homeostatic astrocytes in the gp120Tg mice. Astrocytes expressing Slc6a1, the astrocytic transporter mediating GABA uptake, were mainly in cluster 1 and 2 in the WT spinal cord mice (**Fig. 6a**); shifted to cluster 5 in the gp120Tg cord (**Fig. 6b**), the shift was diminished by Gal3 KO in the gp120TgGal3^-/-^ cord (**Fig. 6c**). As cluster 5 expressing high level of Gfap (**S. Fig. 2b**), the shift of Slc6a1+ astrocytes to cluster 5 in the gp120Tg cord suggesting the activation of these homeostatic astrocytes and the activation were markedly attenuated by Gal3 KO. Same for the astrocytes expressing Slc7a10, the astrocytic transporter mediating excitatory glutamatergic neurotransmission. Astrocytes expressing Slc7a10 were mainly in cluster 1 and 2 in the WT spinal cord mice (**Fig. 6e**); shifted to cluster 5 in the gp120Tg cord (**Fig. 6f**); the shift was prevented by Gal3 KO in the gp120TgGal3^-/-^ cord (**Fig.6g**). Gal3 KO in the WT mice did not impact on these homeostatic astrocytes (**Fig. 6d, 6h**) compared the WT astrocytes (**Fig. 6a, 6e**). Taken together, these data suggest Gal3 KO in the gp120Tg mice attenuated the activation of astrocytes in regulation of neurotransmitter homeostasis.

### Gal3 KO attenuated pathogenic processes of gp120-induced pain

As spinal reactive astrocytes play a key role in gp120-induced pain^7^, we were interested in the potential contribution of the HIPA subtype of reactive astrocytes to the pain pathogenesis. To this end, we first determined the effect of Gal3 KO on the expression of mechanical allodynia in the gp120Tg mice. As shown by previous studies, young adult gp120Tg mice (2 months) do not manifest mechanical allodynia measured by von Frey testing^26^. Nonetheless, we observed that they started to express mechanical allodynia at late ages (4-5 months) (**S**. **Fig. 1**). Importantly, the expression of mechanical allodynia was blocked in gp120Tg/Gal3^-/-^ mice (**Fig. 7a**). These findings indicate that Gal3 is critical for the development of mechanical allodynia in gp120Tg mice.

**Figure 7.**
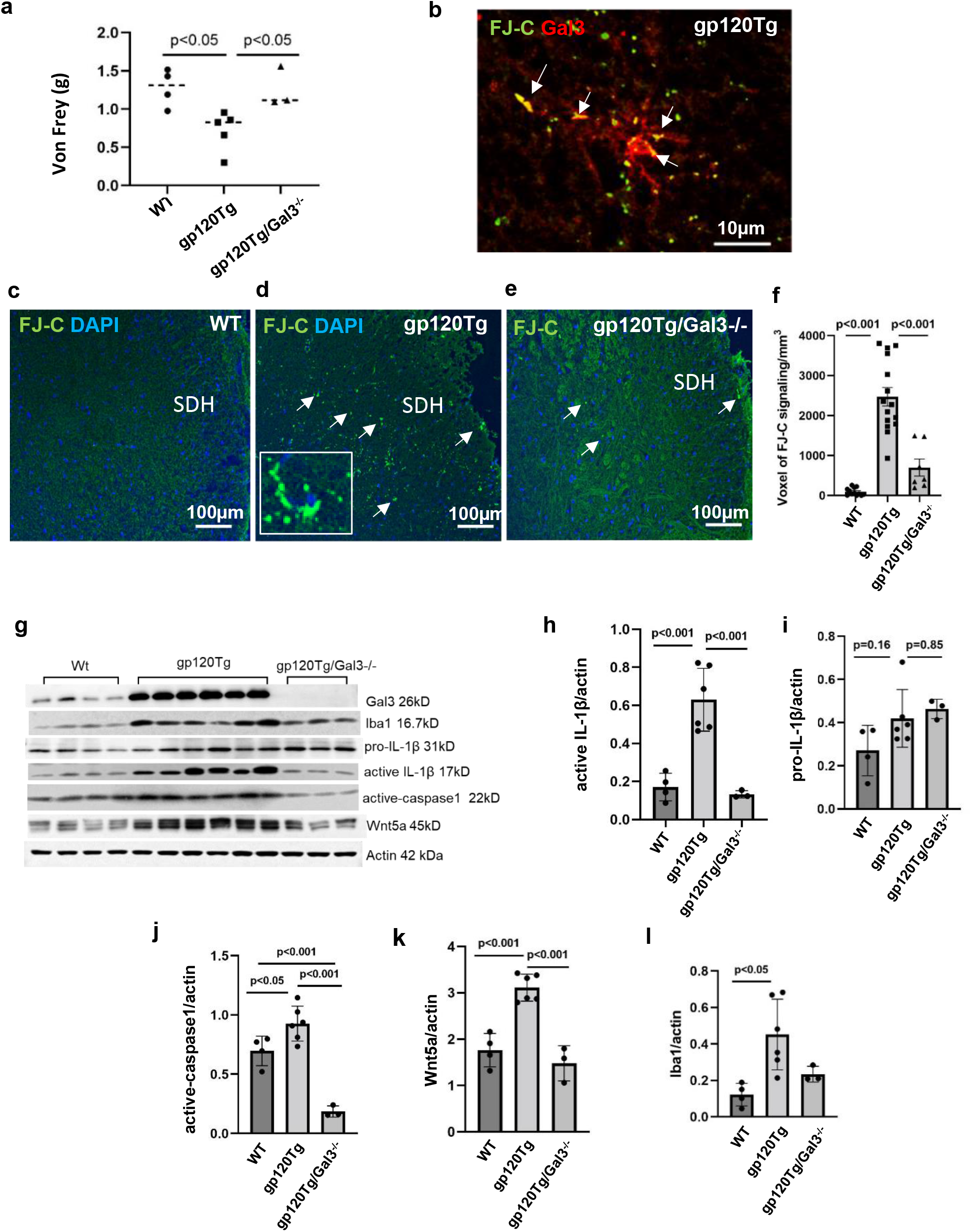
Gal3 KO attenuated mechanical hyperalgesia, neuronal degeneration, and pain related signaling in gp120Tg mice. **a**. Von Frey test of mechanical hyperalgesia. Gp120Tg mice (5 months) developed mechanical hyperalgesia. Gal3 KO blocked the expression of mechanical hyperalgesia of gp120Tg mice. **b.** Association of HIPAs with degenerative neurons in gp120Tg spinal cords. Combination of FJ-C staining and Gal3 IF staining revealed that extensive overlap (arrows) of signals of degenerative neurons (FJ-C staining) and HIPAs (Gal3 IF staining). **c-e.** FJ-C staining of lumbar spinal dorsal horns (SDH) from WT (c), gp120Tg (d), gp120Tg/Gal3-/- (e) mice. FJ-C staining signals (arrows) were markedly increased in gp120Tg mice (d), which were attenuated by Gal3 KO (e). **f**. Quantification of FJ-C staining signals. **g**. Comparison of spinal protein levels of selected molecular markers of neuroinflammation in WT, gp120Tg, gp120Tg/Gal3^-/-^ mice. Protein lysates of lumbar spinal cords were analyzed by immunoblotting of western. **h-i**. Quantification of active Il-1β (h), pro-IL-1β (i), active-caspase1 (j), Wnt5a (k), and Iba1 (l).

Because reactive astrocytes are critical for gp120-induced pain^7^ and Gla3 is critical for the formation of HIPA (**Fig.5e-5g** and **5i**), it is probable that Gal3 controls the expression of gp120-induced allodynia via HIPA. How might HIPA contribute to the pain pathogenesis initiated by gp120? Previous studies on HIV patients autopsies and animal models suggest that neurodegeneration, as indicated by synaptic degeneration, is associated with HIV pain pathogenesis^6,27,28^. As HIPAs are neuronal and synapse phagocytes (**Fig. 4**) and may contribute to the pain pathogenesis via neurodegeneration, we determined the effect of Gal3 KO on neurodegeneration in the SDH. Fluoro-Jade C (FJC) staining was performed to visualize spinal degenerative neurons in WT, gp120Tg and gp120Tg/Gal3^-/-^ mice (5 months). Compared with WT spinal cord, FJC staining signals in gp120Tg spinal cords were significantly increased(**Fig.7c, 7d, 7f**), indicating that gp120 induced neurodegeneration. We also observed that FJC signals co-localized with HIPA (**Fig.7b**), suggesting that HIPAs were closely associated with degenerative neurons. Importantly, Gal3 KO blocked the increase of FJC signals in spinal cords of gp120Tg mice (**Fig. 7e, 7f**).

The Neuroinflammation in the SDH is a specific biomarker for pain development in HIV patients^6^. In particular, activation of astrocytes is a hallmark of neuroinflammation in the SDH specifically from the HIV patients with chronic pain^2^. These previous findings provide strong premise for the hypothesis that HIPAs are among the reactive astrocytes that contribute to the pathogenic neuroinflammation underlying HIV-associated pain. Data from snRNA-seq analysis reveals upregulated expression of numerous inflammatory genes (**Fig. 1k**) (**S. Fig. 3**) in HIPAs. To test this hypothesis, we compared protein levels of various markers of neuroinflammation in the spinal cords of WT, gp120Tg, and gp120Tg/Gal3^-/-^ mice (5 months) by immunoblotting analysis (**Fig.7g**). We chose to focus on neuroinflammation markers IL-1β, Wnt5a and reactive microglia because of their confirmed contribution to the development of gp120-induced pain^7^.

We observed that cleaved IL-1β was upregulated in gp120Tg spinal cords, but this upregulation was abolished in gp120Tg/Gal3^-/-^ mice (**Fig.7h**). On the other hand, the levels of pro-IL-1β were not significantly different among WT, gp120Tg and gp120Tg/Gal3^-/-^ mice (**Fig.7i**). Because reactive astrocytes are the source of gp120-induced IL-1β^7^ and Gal3 KO diminished HIPA (**Fig. 5g, 5i**), these data indicate that Gal3 likely controls IL-1β in HIPAs Further, these data also suggest that Gla3 regulates IL-1β at the stage of its activation/processing rather than transcription and translation. To gain insight into the potential Gal3-regulated IL-1β activation mechanism, we tested the involvement of inflammasome. We measured the levels of active-caspase 1, which is the inflammasome-associated pro-IL-1β processing proteinase and observed that active-caspase1 was significantly upregulated in gp120Tg mice (**Fig.8j**). Strikingly, Gal3 KO not only abolished the upregulation but also brought the active-caspase1 level drastically below the basal level (**Fig. 7j**). These results indicate that Gal3 regulates gp120-induced IL-1β via controlling the activation of caspase1.

**Figure 8.**
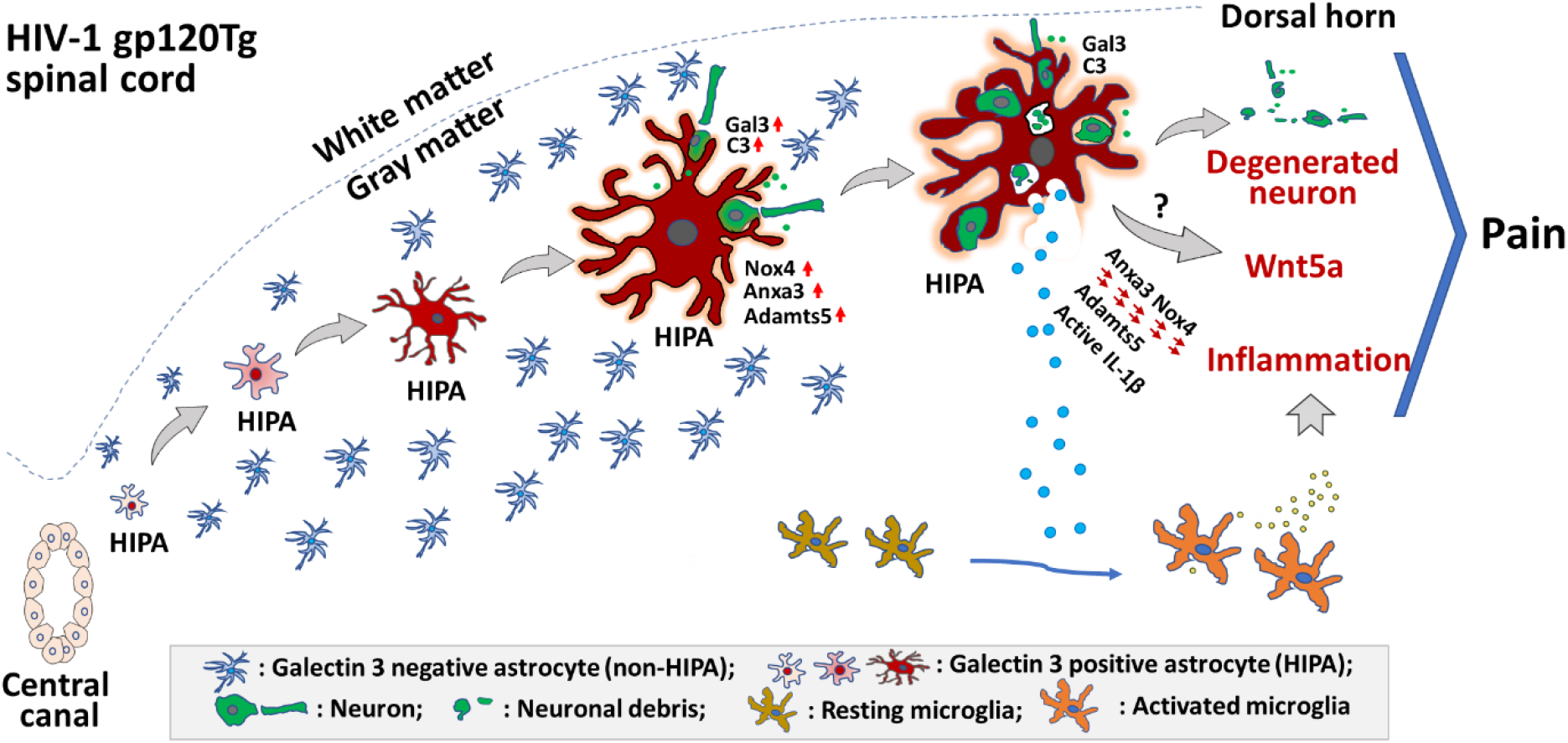
A model of HIPA differentiation in the spinal cord of gp120Tg mice. HIPAs are a novel subpopulation of reactive astrocytes, with the expression of phagocytosis-related genes (Gal3, C3 etc.) and inflammation-related genes (Anxa3, Nox4, Adamts5 etc.), in the lumbar spinal cord of gp120Tg mice. HIPAs originate from the Gal3^+^ stem cells lined in spinal central canal, migrating along a clear trajectory toward the spinal marginal area with gradual increase of cell volume and complexity. During this process, Gal3 protein is progressively translocated from the nucleus to the cytoplasm and plasma membrane. HIPAs are eventually developed into enormous, big phagocytes. Gal3 (and probably other proteins such as C3) released from HIPAs may opsonize nearby neurons and synapses for phagocytosis. HIPAs also express inflammatory proteins such as Anax3, Nox4 and Adamts5. The upregulation of caspase1 in HIPAs may lead to the activation of inflammasome pathway that activate IL-1β. These inflammatory cytokines released from HIPAs may activate microglia, which further elevate spinal inflammation. Via an unknown mechanism, HIPAs may also regulate the expression of Wnt5a, a key protein that regulates neuroinflammation during the pathogenesis of gp120-induced pain. Both the phagocytotic and inflammatory activities of HIPAs may contribute to the damages of neural circuits in the spinal dorsal horn and the expression of pathological pain induced by HIV-1 gp120.

Wn5a is a critical neuroinflammation regulator that is specifically upregulated in the SDH of pain-positive HIV patients^29^ and plays a key role in gp120-induced pain pathogenesis^7,30^. We found that spinal Wnt5a was significantly upregulated in gp120Tg mice and Gal3 KO significantly attenuated this upregulation in gp120Tg/Gal3^-/-^ mice (**Fig.7k**).

Reactive microglia are a major source of inflammatory mediators during the development of neuroinflammation and are critical for the expression of the early phase of gp120-induced pain^7^. Consistent with results from previous studies^27^, we observed upregulation of microglia/macrophage marker Iba1 in the spinal cord of gp120Tg mice (**Fig.7l**), indicating the activation of microglia. Furthermore, we found that Gal3 knockout attenuated the Iba1 upregulation (**Fig.7l**) indicating an inhibitory effect of Gal3 KO on microglial activation. Collectively, the above analyses on multiple markers suggest that Gal3 plays key roles in controlling the formation of HIPAs and in establishing neuroinflammation critical for the development of gp120-induced pain.

## Discussion

By snRNA-seq, we identify a novel astrocyte subtype, HIPAs, from the lumbar spinal cord of HIV gp120Tg mice. HIPAs express Gal3, possess transcriptomic signatures of inflammation and phagocytosis, and phagocytose neuronal components. HIPAs are depleted after Gal3 KO. Concomitantly, neuroinflammation and neurodegeneration are attenuated, and the expression of mechanical allodynia is blocked. The data collectively suggest HIPAs may contribute to pathogenesis of HIV-associated pain by promoting neuroinflammation and neurodegeneration in the spinal pain neural circuits.

### A novel subtype of reactive astrocytes in pain pathogenesis

Reactive astrocytes emerge as a key contributor to the development of pathological pain^4,5^. The role of reactive astrocytes in the pathogenesis of HIV-associated pain was suggested by the observations of astrogliosis specifically in the SDH of HIV patients with pain^2^ and blockage of pain development after ablation of reactive astrocytes in mouse models^7^. However, the specific subtype of astrocytes underlying the pathogenesis of HIV-associated pain was unclear. We found HIPAs were induced in the spinal cord of gp120Tg mice, and their depletion by Gal3 KO was associated with diminished expression of allodynia. Hence, HIPAs are a specific subtype of reactive astrocytes associated with HIV-related pain pathogenesis in animal models.

### Potential pathogenic mechanism mediated by the HIPAs

HIPAs express multiple upregulated inflammatory genes, including Annexin A3 (Anxa3), Nox4, and Adamts5. Upregulations of these genes are implicated in pain pathogenesis in various neuropathic pain models^31,32,33^. Anxa3 was upregulated in the spinal cords of a rat neuropathic pain model induced by chronic constriction injury (CCI), and its inhibition alleviated CCI-induced mechanical allodynia and thermal hyperalgesia^31^. Nox4, one of the major the source of intracellular reactive oxygen species (ROS)^34^, promoted caspase 1 production and IL-1β activation^35^; both ROS and IL-1β are known to play important roles in pain pathogenesis^36,37^. ADAMTSs are a family of zinc metalloendopeptidases that participate in diverse biological processes including extracellular matrix remodeling and inflammation^38^. Adamts5 is a protease of Reelin, the extracellular matrix glycoprotein in the spinal cord dorsal horn^39^. Reduction of Reelin resulted in mechanical and thermal hyperalgesia^33,40^. The upregulated expression of these inflammatory genes in HIPAs suggest HIPAs are inflammatory cells that contribute to pain pathogenesis. Consistent with this notion, HIPA depletion induced by Gal3 KO significantly attenuated the expression of these genes (**Fig.5j-5l, 5n-5p**) and neuroinflammation in the spinal cords of gp120Tg/Gal3 KO mice, as shown by the reversal of upregulated proinflammatory mediator IL-1β (**Fig. 7h**) and proinflammatory regulator Wnt5a (**Fig. 7k**), both of which are critical for gp120-induced pain pathogenesis^7,30^.

HIPAs also express high levels of phagocytosis-related genes, including Lgals3, C3, Palld. Gal3 (Lgals3) and C3 are phagocytosis ligand. Deposition of Gal3 and C3 on the surface of target cells opsonizes the target cells and drives phagocytosis^41,42^. Anxa3, an inflammation-related gene expressed in HIPAs, also plays a specialized role in phagocytosis, by mediating calcium-dependent granules-phagosome fusion^43^. The expression of phagocytosis-related genes suggest HIPAs are phagocytes. Indeed, we observed phagocytosed somatic and synaptic components in HIPAs (**Fig.4**). In addition, depletion of HIPAs is associated with significant decrease of neuronal degeneration in gp120Tg/Gal3 KO mice (**Fig. 7f**). Collectively, these results suggest HIPAs may contribute to neurodegeneration (including synapse loss) in the pain neural circuits. Previous studies show that microglia also play a critical role in the synaptic degeneration^27^. It will be interesting for future studies to determine how HIPAs and microglia coordinate gp120-induced synapse degeneration in the spinal neural circuits. The relevance of synaptic degeneration in pain pathogenesis of HIV-associated pain is suggested by the finding that synapse loss is specifically observed in the SDH of HIV patients who developed pathological pain but not in HIV patients without pain complication^44^.

### HIPAs in regulation of neurotransmitter homeostasis

A fundamental function of astrocytes is to maintain neurotransmitter homeostasis in extracellular space, by transporter-mediated neurotransmitter uptake and gliotransmitter release^45^. During astrogliosis, astrocytes undergo a spectrum of biological changes that involve proliferation, morphological alterations, and changes of cellular functions^46^. Our snRNA-seq data indicated that the total number of astrocytes in the gp120Tg spinal cord did not increase (**S. Fig.7**). Instead, the astrocytes manifested a marked shift from the GFAP^low^ population to GFAP^high^ population (**S. Fig. 2**). GFAP^low^ astrocytes show high expression levels of Slc6a11 which regulates gamma-aminobutyric acid (GABA) uptake^18^ and Slc7a10 which is a sodium-independent amino acid transporter with a primary role related to modulation of excitatory glutamatergic neurotransmission^19^. Concomitantly, Slc6a11^+^ and Slc7a10^+^ astrocytes were shifted from GFAP^low^ population (**Fig. 6a, 6e, S. Fig. 2a**) to GFAP^high^ population in the gp120Tg spinal cord (**Fig. 6b, 6f**, **S. Fig. 2b**) suggesting an activation of the homeostatic astrocytes which is expected to be associated with an impairment in neurotransmitter homeostasis in the pain neural circuits. Depletion of HIPAs markedly diminish the activation of these homeostatic astrocytes in the gp120 spinal cords (**Fig. 6c, 6g**).

HIPAs express marker genes related to inflammation and phagocytosis. HIPAs do not express astrocytic transporter genes Slc6a11 and Scl7a10. This transcriptomic signature suggests HIPAs have lost the capacity of regulating neurotransmitter homeostatic function.

### Gal3 in regulation of HIPA astrogliosis

HIPA were almost completely depleted in the spinal cord of gp120Tg/Gal3^-/-^ mice (**Fig. 5g, 5i**). This observation suggests a key role of Gal3 in HIPA genesis. In the WT mice, Gal3 was expressed at a low basal level in cells restricted in regions around the spinal central canal (**Fig.3b, 3e, 3h**). In gp120Tg mice, Gal3 protein expression was markedly upregulated (**Fig.3a**), but Gal3^+^ cells in the central canal were diminished (**Fig.3c, 3i, 3l**). Instead, Gal3^+^ cells migrated out of the central canal region (**Fig. 3i, 3l**), forming a trajectory from the central canal to the spinal marginal area in the SDH (**Fig. 3d**). This is consistent with the idea that HIPAs are originated from the Gal3^+^ cells in the central canal lined with ependymal cells, which are a pool of neural stem cells with the potential of differentiation into discrete multilineages^47^. Previous studies indicate that Gal3 plays a key role in glial fate choices in stem cell differentiation in the postnatal subventricular zone of mouse brains^48^. Overexpression of Gal3 promotes astrocytic differentiation, and simultaneously suppresses oligodendrogenesis^48^ . Indeed, compared to the Gal3^+^ cells in the WT central canal which did not express Gfap, the expression of Gfap in the gp120Tg central canal cells suggested the central canal cells showed astrocytic lineages in the gp120 mice. Our data, together with the previous findings, indicate that Gal3 controls HIPA differentiation from the stem cells in the spinal central canal in gp120Tg mice.

In addition to controlling HIPA genesis, Gal3 is probably also critical for HIPAs to develop into functional phagocytes, because of its role in opsonizing target cells. As HIPAs reach the dorsal horn, the cellular distribution pattern of Gal3 protein expands (from the nucleus) to cytoplasm and plasma membranes. At this stage, HIPAs become phagocytes of neuronal and synaptic components (**Fig. 4**), with increased cell volume and plasma Gal3. These mature HIPA phagocytes may release Gal3 to opsonize neurons for phagocytosis. It is suggested that Gal3 in extracellular space interacts with plasmatic membrane receptors by binding to both glycosylated cargos and membrane glycosphingolipids on cell surface^49^ and induces inward membrane bending and initiation of endocytosis in a clathrin-independent manner^50^. Whether this mechanism apply for the phagocytosis of HIPAs needs further investigation.

In summary (**Fig. 8**), HIPAs are a novel subtype of reactive astrocytes induced in the gp120Tg spinal cord. They show characteristic expression of inflammation- and phagocytosis-related genes. Based on our data, we propose the following model. HIPAs originate from the Gal3+ cells from the spinal central canal and migrate to spinal marginal areas. During this process, HIPAs gradually increase their cell volume and process complexity, expand the cellular distribution pattern of Gal3 protein from nucleus to the cytoplasm and plasma membrane, and eventually mature into functional phagocytes. The released Gal3 and probably other proteins such as C3 from phagocytotic HIPAs opsonizes nearby neurons and synapses for phagocytosis. HIPA-mediated phagocytosis in spinal dorsal horn may result in neuronal damage. HIPAs also express inflammatory proteins that are pain mediators, including Anax3, Nox4 and Adamts5. These inflammatory cytokines released from HIPAs may activate microglia, which further elevate spinal inflammation. Via an unknown mechanism, HIPAs may also regulate the expression of Wnt5a, a key protein that regulates neuroinflammation during the pathogenesis of gp120-induced pain. These findings suggest that HIPAs may contribute to gp120-induced pain pathogenesis by phagocytotic and inflammatory pathways.

## Method

### Animal

Gp120 Tg mice (from Dr. Marcus Kaul, Sanford-Burnham Medical Research Institute) express HIV-1 LAV gp120 under the control of the glial fibrillary acidic protein (Gfap) promoter^15^. Lgals3 Homozygotes knockout (Gal3^-/-^) mice were purchased from the Jackson Laboratory (Strain: #006338). All animal procedures were performed according to protocol 0904031B approved by the Institutional Animal Care and Use Committee at the University of Texas Medical Branch, and all methods were performed in accordance with the relevant guidelines and regulations.

### Spinal cord lumbar dissection

We used the same tissue dissociation procedures to prepare spinal nuclei dissociation from all groups in parallel. To isolate nuclei for snRNA-seq, the WT, gp120Tg, gp120Tg/Gal3^-/-^, Gal3^-/-^ mice were euthanized with 14% urethane followed by decapitation. The lumbar spinal cords were rapidly dissected out and temporarily stored in 6ml cold Hibernate A/B27 (HABG) medium for further process. HABG was prepared by adding 30 ml of Hibernate A (BrainBits, cat. no. HA), 600μl of B27 (ThermoFisher, cat. no.17504044), 88 μl of 0.5mM Glutamax (Invitrogen, cat. no. 35050-061), penicillin-streptomycin (ThermoFisher, cat. no.15070063) to final concentration of 2%, and DNase I (ThermoFisher, cat. no. AM2224) to final concentration of 80 U/ml.

### Nucleus dissociation from the spinal cord lumbar

Three lumbar spinal sections from each group were pooled for nuclei dissociation. Nuclei were dissociated following the detergent-mechanical cell lysis method^51^. Briefly, placed the spinal lumbar sections in a pre-chilled Dounce homogenizer. Added 500 μl pre-chilled detergent lysis buffer. The detergent lysis buffer was prepared by adding 3μl 20% Triton-X to 1800 μl low sucrose buffer (Sucrose: 0.32 M; HEPES (pH = 8.0):10mM; CaCl2: 5mM; MgAc: 3 mM; EDTA: 0.1mM; DTT: 1mM). Dounce with 5 strokes of pestle A (‘loose’ pestle), then 5-10 strokes of pestle B (‘tight’ pestle). Avoided lifting the homogenizer out of the lysis solution in between strokes and avoided introducing bubbles. Placed a 40 μm strainer over a pre-chilled 50 ml conical tube and prewet with 1 ml of low sucrose buffer. Added 1 ml of low sucrose buffer to the Dounce homogenizer containing the crude nuclei in the lysis buffer and mixed gently by pipetting 2–3 times. Passed the crude nuclei prep over the 40 μm strainer into the pre-chilled 50 ml conical tube. Passed an additional 1 ml low sucrose buffer over the 40 μm strainer, bringing the final volume to 3 ml of the low sucrose buffer and 500 μl of the lysis buffer.

Centrifuged the sample at 3,200 × g for 10 min at 4 °C. Decanted the supernatant. Resuspended the pellet using 3 ml of low sucrose buffer. Gently swirled to remove the pellet from the wall to facilitate the resuspension. Let the sample sit on ice for 2 min and transferred the suspension to an Oak Ridge tube. Using the homogenizer (IKA, Ultra Turrax T8) at setting 1, homogenized the nuclei in low sucrose buffer for 15–30 s, keeping the sample on ice. Using a serological pipette, layered 12.5 ml of density sucrose buffer (Sucrose: 1M; HEPES (pH=8.0): 3mM; MgAc: 3mM; DTT: 1mM) underneath the low sucrose buffer homogenate, taking care not to create a bubble that disrupts the density layers. Centrifuged the tubes at 3,200 × g for 20 min at 4 °C. Once the centrifugation was complete, immediately decanted the supernatant in a flicking motion. Using 500 μl of resuspension solution, resuspended the nuclei remaining on the wall. Filtered the nuclei through a 30–35 μm pore-size strainer and collect in a pre-chilled tube. Determined the nuclei yield using a hemacytometer to count nuclei under a 10X objective. Proceeded 10x single-nucleus RNA-sequencing (snRNA-seq).

### Generation snRNA-seq library and feature-barcode matrices

Cell suspensions were loaded into the 10X Genomics Chromium single-cell microfluidics device with the Single Cell 3’ Library & Gel Bead Kit v3 (10X Genomics, cat. no. PN-1000075) according to the manufacturer’s instructions. 12 PCR cycles were used for cDNA amplification, and 13 to 15 PCR cycles were performed for ligating adaptors. The library quality and size were checked using Bioanalyzer High Sensitivity DNA Analysis (Agilent Technologies, USA). The libraries were sequenced in Single Cell Genomics Core, College Baylor College of Medicine, USA.

### Seurat snRNA-seq integration analysis

Raw sequencing data were pre-processed with Cell Ranger (v.3.1.0) (10X Genomics). The generated filtered single-cell expression matrices files from the WT, gp120Tg, gp120Tg/Gal3^-/-^, Gal3^-/-^ mice were imported into R for Seurat snRNA-seq integration analysis. The Seurat R package (v.4.0.5) was used for dimension reduction and cluster analysis. By default, Seurat implements a global-scaling normalization method “LogNormalize” that normalizes the gene expression measurements for each nucleus by the total expression, multiplies this by a scale factor (10,000 by default), and log-transforms the result. We set the filter criteria of nFeature_RNA > 200 and nFeature_RNA < 5000 as a QC control for snRNA-seq data to remove the dead, low-quality nuclei, and doublets. This was followed by normalization prior to selecting all highly variable genes falling within a selected cut-off window for PCA clustering. Statistically significant PCs (p-value < 0.001) were used in cluster determination. Uniform Manifold Approximation and Projection (UMAP) plots were produced at a resolution of 0.6. Clusters were annotated based on the expression level of canonical marker genes and gene expression visualized using feature maps. Cluster of astrocytes were identified using the classic marker gene GFAP, Aqp4, Gja1. Astrocyte clusters from the integrated UMAP plot were subset for astrocyte subset analysis. UMAP plots of astrocyte were produced at a resolution of 0.5. The integrated UMAP plot were split by the genotypes to view the Astrocyte cluster at each condition.

### Immunofluorescent (IF) staining

Mice were anesthetized with euthanized with 14% urethane and transcardially perfused with 30 ml cold 1×PBS. The spinal cords L4–L5 lumbar spinal cord segments were collected, and fixed in freshly prepared 4% paraformaldehyde for 24 h at 4°C. The tissues were then immersed in 30% sucrose in 1xPBS at 4°C until they sank to the bottom, embedded in OCT mounting medium, and stored in an air-tight container at −80°C prior to sectioning. The mounted tissue blocks were equilibrated to −20°C in a cryostat for ~1 hour prior to sectioning. 10–35 μm cryo-sections were cut, mounted onto SuperFrost® Plus slides or stored in Hito floating section storage solution (Hitobiotec) at −20°C until they were stained for immunocytochemistry. For immunostaining, sections were rinsed with 1xPBS two times to remove storage solution and blocked with 5% BSA and 0.3% Triton X-100 in 1xPBS for 2 h at room temperature. This was followed by 48 h incubation with primary antibodies. The primary antibodies used included anti-Gal3 antibody (1:500, Santa Cruz, cat. no. sc-23938), anti-GFAP antibody (1:500, Millipore, cat. No. AB5541), anti-Iba1 antibody (1:500, 1:1000; Wako, cat. No. 016–20001), ant-Vimentin antibody (1:500, abcam, cat. no. ab137321), anti-NeuN antibody (1:500, Millipore, cat. no. ABN90), and anti-vGlut1 antibody (Synaptic systems, cat. no. 35 303). After five washes with PBS, the sections were incubated with the corresponding second antibodies at room for 2h. The stained sections were mounted with ProLong Gold Antifade Mountant (Invitrogen, cat. no. P36930) and stored overnight at 4°C. Images were viewed using Zeiss LSM 880 with Airyscan (Carl Zeiss Microscopy GmbH) and Nikon A1R MP (Nikon Instruments Inc.). The 3-D images were processed using Imaris 9.6 software (Oxford Instruments).

### Fluoro-Jade C (FJC) staining

10μm cryo-sections of spinal lumbar sections were prepared for FJC staining. FJC staining was performed using Fluoro-Jade C Ready-to-Dilute Staining Kit for identifying degenerating neurons (Biosensis, cat. no. TR-100-FJ) according to manufacturer’s recommendation. After the FJC staining, IF staining was performed on the same sections using the anti-Gal3 antibody (1:500, Santa Cruz, cat. no. sc-23938) according to the protocol described above. The stained sections were mounted with ProLong Gold Antifade Mountant (Invitrogen, cat. no. P36930) and stored overnight at 4°C. Images were viewed using Zeiss LSM 880 with Airyscan (Carl Zeiss Microscopy GmbH) and Nikon A1R MP (Nikon Instruments Inc.).

### Western blotting analysis

Mice were killed with excess anesthesia (isoflurane), and the L4–L5 lumbar spinal cord segments of mice were collected and homogenized in RIPA lysis buffer (50 mm Tris-HCl, pH 7.4, 150 mm NaCl; 0.1% SDS, 1% Nonidet P-40; 10% glycerol, 1 mm EDTA, pH 8.0, containing PMSF and protease inhibitor mixture (Sigma-Aldrich, cat. no. #P8340). Protein concentration in the homogenates was determined using a Pierce BCA Protein Assay Kit (Thermo Fisher Scientific, cat. no. 23227). Protein concentration was titrated to 2 μg/μl for immunoblotting analysis. Equal amounts of protein (30 μg/lane) were loaded and separated by SDS-PAGE, followed by transferring to polyvinylidene difluoride membranes (Millipore, cat. no. IPVH00010). Immunoblots were blocked by 5% nonfat milk in Tris-buffered saline–Tween-20 (20 mm Tris-HCl, 150 mm NaCl, pH 7.5, 0.1% Tween-20) for 2 h at room temperature. The blots were then sequentially incubated with primary and secondary antibodies. Primary antibodies included anti-Gal3 (1:500, Santa Cruz, cat. no. sc-23938); anti-cleaved Il1β (1:500, Millipore, cat. no. ab1413-I); anti-pro-Il1β (1:500, Cell Signaling Technology, cat. no. #12242); anti-cleaved caspase-1 (1:1000, Cell Signaling Technology, cat. no. #4199) and anti-wnt5a (1:1000, Abcam, cat. no. ab72583); anti-Iba1 (1:1000; Wako; cat. no. AB_839506); anti-actin (1:1000; Santa Cruz, cat. AB_630836). Protein bands were visualized using the Pierce Enhanced Chemiluminescence Kit (Thermo Fisher Scientific, cat. no. 32106). β-Actin was blotted as a loading control. The intensity of bands was quantified by densitometry analysis with NIH ImageJ.

### Mechanical pain behavioural test

Paw withdrawal thresholds (PWTs) were measured by von Frey testing on the plantar surface of the hind paw. Three days before testing, mice were habituated to the testing surroundings for 2 h per day. On the testing day, a series of calibrated von Frey filaments (0.1 to 2.0 g) were applied perpendicularly to the central area of the hind paw. The PWT values were calculated using the Dixon “up and down paradigm” method. To minimize the subjective effect and bias, behavioral tests were performed under double-blind conditions. The experimenter did not know the genotype of individual animals.

## Notes

### Competing Interest Statement

The authors have declared no competing interest.

### Summary of Updates

1. Replaced Figure 3d with an image which shows a better HIPA trajectory from the central canal to SDH. 2. Moved Figure 5 (Western of Gfap and Vim) to the Supplementary section (S. Figure 6). 3. Added Figure 3e-3i, the high magnifications of Gal3+ cells in the WT and gp120Tg central canals. 4. Added Figure 5n-5u in support of the diminish of HIPAs by Gal3 KO. 5. Added Figures 6a-6h to support the point in the abstract and discussion that Gal3 KO in the gp120Tg mice attenuated the activation of astrocytes in regulation of neurotransmitter homeostasis. 6. Revised the summary figure (Figure 8). 7. Deleted the unnecessary supplementary tables (S. table 3- S. tables 7). Using supplementary figures (S. Figure 3, S. Figure 4 and S. Figure 7) instead.

