## Supplemental Figures for "A novel subtype of reactive astrocytes critical for HIV associated pain pathogenesis"

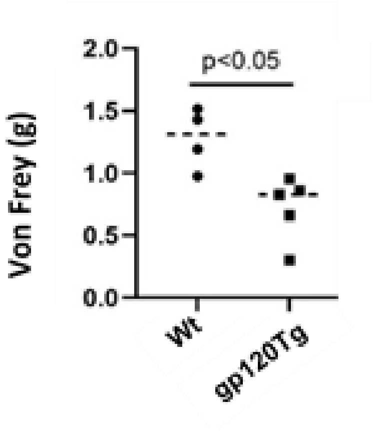


**Supplementary Figure 1**. Von Frey test demonstrated gp120Tg mice developed mechanical allodynia at late ages (4-5 months)

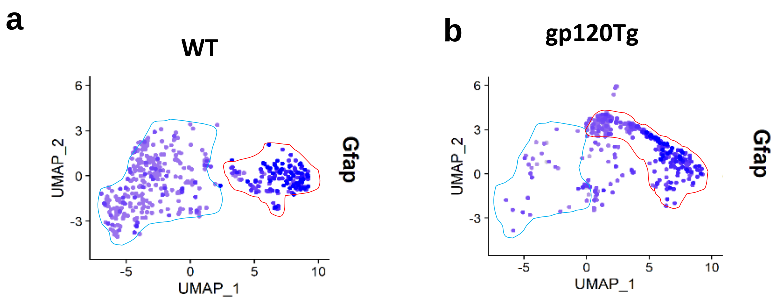


**Supplementary Figure 2.** UMAP plots demonstrate the shift of astrocytic clusters in the gp120Tg mice. **a**, UMAP demonstrated the astrocytic clusters expressing low level of Gfap (Gfap$^{low}$) (surround by blue line) and high level of Gfap (Gfap$^{high}$) (surround by red line) in the WT mice. **b**, The astrocytic clusters in the gp120Tg mice manifested a clear shift from Gfap$^{low}$ toward Gfap$^{high}$ populations. Dark blue dots, high Gfap expression, light blue dots: low Gfap expression.

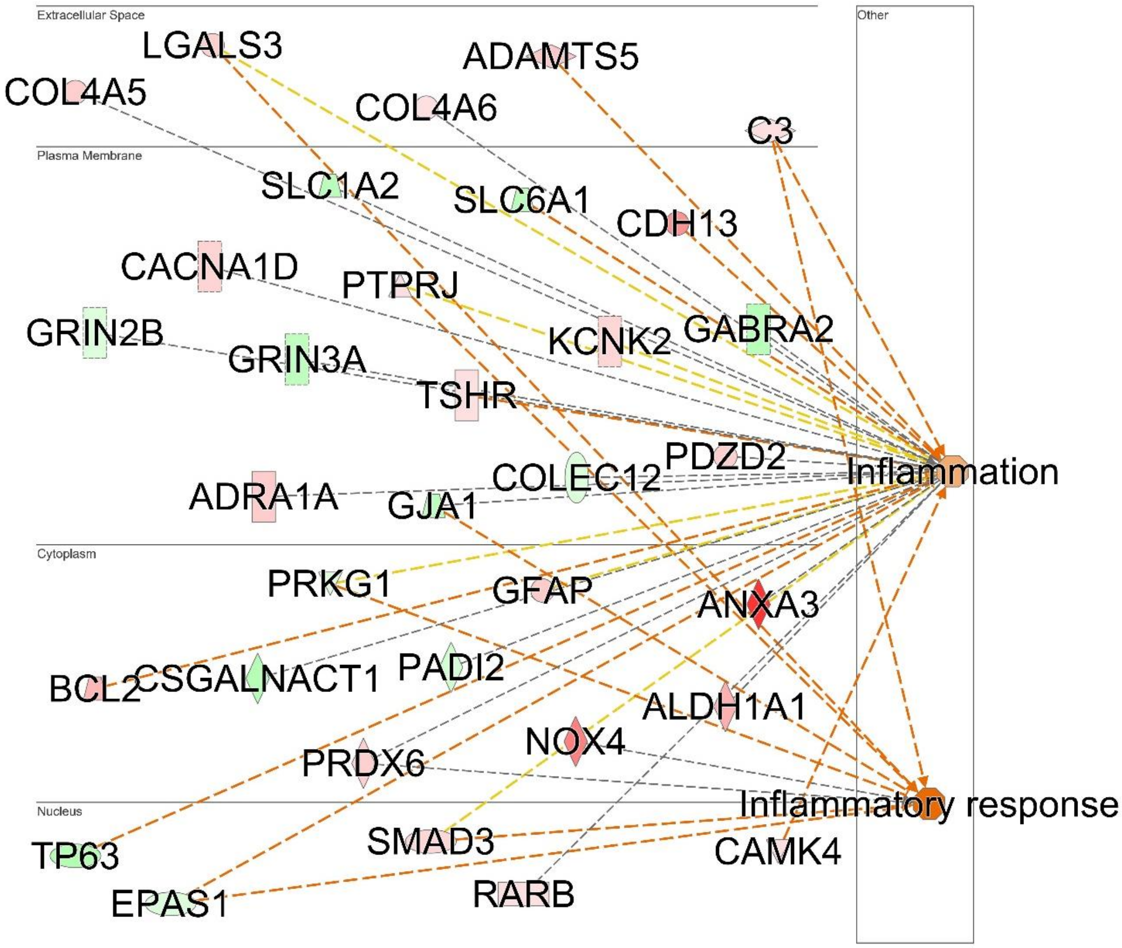


**Supplementary Figure 3.** Ingenuity Pathway Analysis (IPA) predicted HIPAs were associated with an increased inflammatory response**.** The 112 HIPA (cluster 7)-specific marker genes with adjusted p-value < 0.05 from the gp120Tg spinal cord were imported into Ingenuity Pathway Analysis (IPA). IPA identified 33 well-characterized inflammatory genes. Green: down-regulated; red: up-regulated; the orange dot indicated increased inflammatory functions of HIPAs.

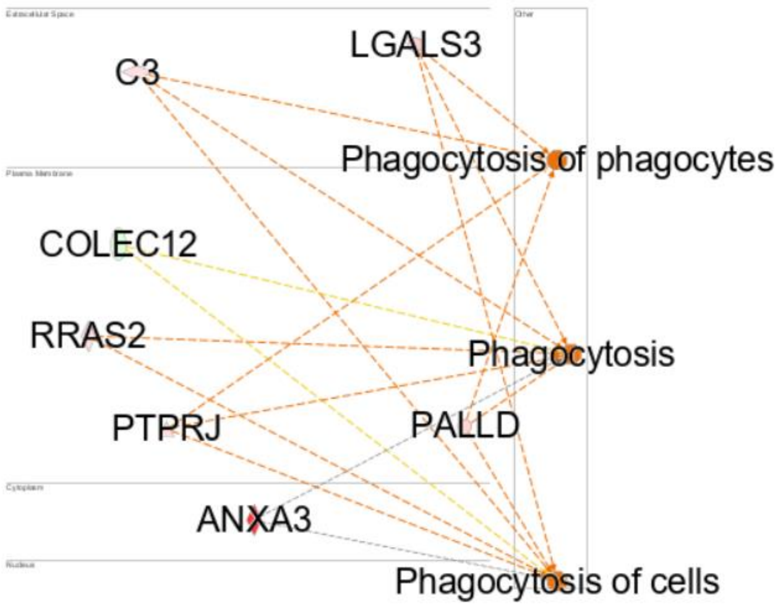


**Supplementary Figure 4.** IPA predicted HIPAs were associated with an increased phagocytotic function. The 112 HIPA (cluster 7)-specific marker genes with adjusted p-value < 0.05 from the gp120Tg dataset were imported into Ingenuity Pathway Analysis (IPA). IPA identified 7 well-characterized inflammatory genes. Green: down-regulated; red: up-regulated; the orange dot indicated increased phagocytotic functions of HIPAs.

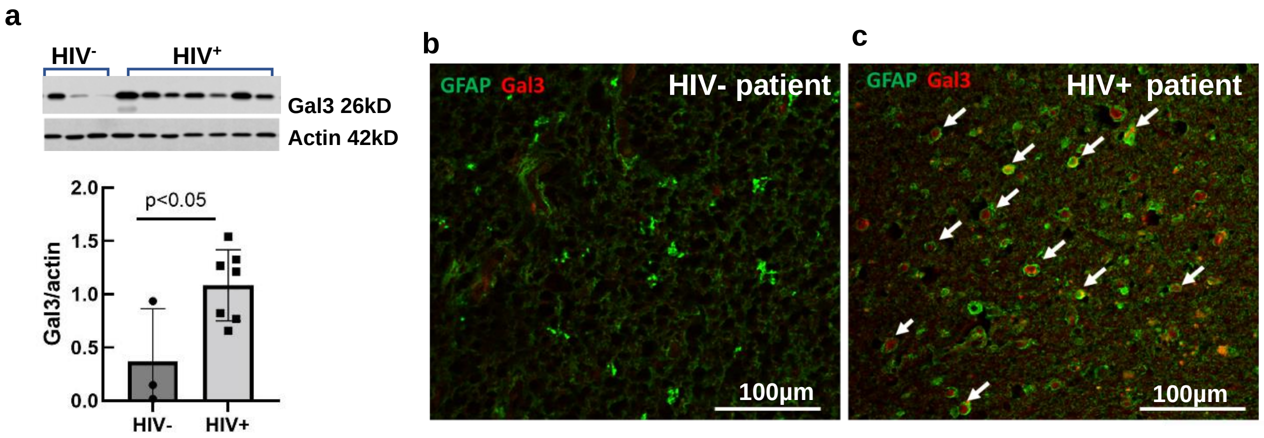


**Supplementary Figure 5.** The upregulation of Galectin-3 (Gal3) protein expression in the spinal cords of human HIV$^+$ patients**. a,** Western blot and quantification demonstrated the significantly increased Gal3 protein expression in spinal cords of the patients died of HIV vs patients died of endocarditis, pulmonary embolus, and pneumonia respectively. **b** and **c**, Immunofluorescence (IF) staining revealed abundant Gal3 (red, white arrows) protein expressed in the spinal cord of a deceased HIV$^+$ patient (**c**) but not in the spinal cord of a patient died from endocarditis (**b**). White arrows point to the expression of Gal3 in astrocytes. Note: the astrocyte shape was severely distorted in the deceased HIV patient for unknow reasons.

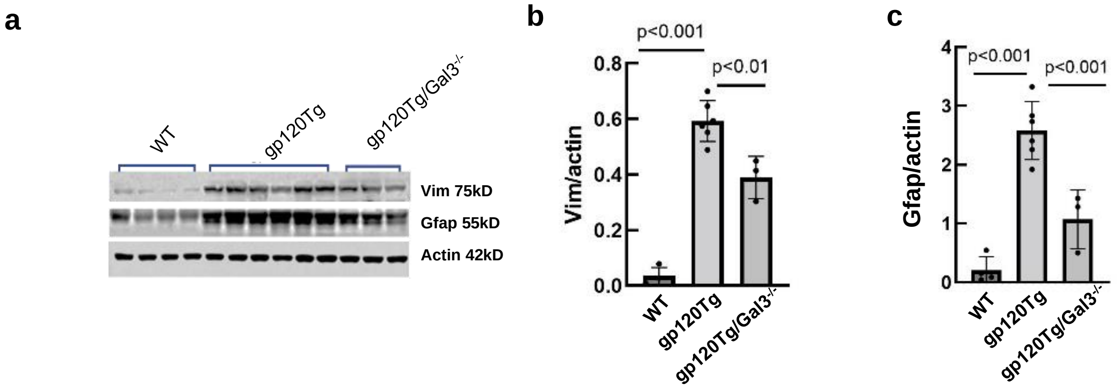


**Supplementary Figure 6.** Gal3 knockout (Gal3KO) in the gp120Tg mice significantly attenuated the increased expression of reactive astrocytic marker proteins Vimentin (Vim) and Gfap in the lumbar spinal cords**. a**, Western blots of Vim and Gfap in the lumbar spinal cord of the WT (n=4), gp120Tg (n=6), gp120Tg/Gal3-/- (n=3) mice; **b** and **c**, Quantification of the protein expression of Vim (b) and Gfap (c).

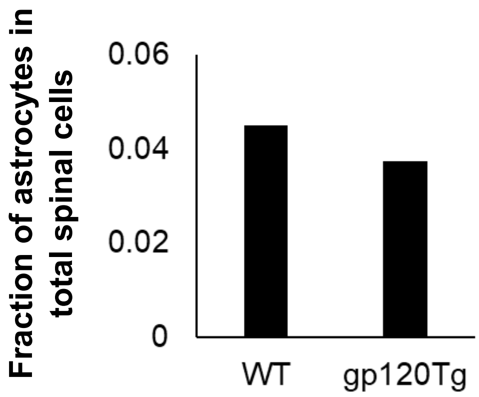


**Supplementary Figure 7.** snRNA-seq revealed the fraction of astrocytes in total lumbar spinal cells from the WT (n=3) and gp120Tg (n=3) mice.
